# Heterocomplexes between the Atypical Chemokine MIF and the CXC-Motif Chemokine CXCL4L1 Regulate Inflammation and Thrombus Formation

**DOI:** 10.1101/2021.11.26.470090

**Authors:** Markus Brandhofer, Adrian Hoffmann, Xavier Blanchet, Elena Siminkovitch, Anne-Katrin Rohlfing, Omar El Bounkari, Jeremy A. Nestele, Alexander Bild, Christos Kontos, Kathleen Hille, Vanessa Rohde, Adrian Fröhlich, Jona Golemi, Ozgun Gokce, Christine Krammer, Patrick Scheiermann, Nikolaos Tsilimparis, Wolfgang E. Kempf, Lars Maegdefessel, Remco T.A. Megens, Hans Ippel, Rory R. Koenen, Kevin H. Mayo, Meinrad Gawaz, Aphrodite Kapurniotu, Christian Weber, Philipp von Hundels-hausen, Jürgen Bernhagen

## Abstract

To fulfil their orchestrating function in immune cell trafficking in homeostasis and disease, a network of 49 chemokines and 23 receptors capitalizes on features of specificity, redundancy, and functional selectivity such as biased agonism. The discovery of the chemokine interactome, i.e. heteromeric chemokine-chemokine interactions, even across CC- and CXC-class borders, has further expanded the complexity within the network. Moreover, some inflammatory mediators, which are not structurally linked to classical CC-, CXC-, CX_3_C-, or C-chemokines, can bind to chemokine receptors and behave as atypical chemokines (ACKs). We identified the cytokine macrophage migration inhibitory factor (MIF) as an ACK that binds to the chemokine receptors CXCR2 and CXCR4 to promote atherogenic leukocyte recruitment. Here, we hypothesized that chemokine-chemokine interactions extend to ACKs and that MIF may form heterocomplexes with classical chemokines. We tested this hypothesis, applying an unbiased chemokine protein binding array. The platelet chemokine CXCL4L1, but not its variant CXCL4 or the CXCR2/CXCR4 ligands CXCL8 or CXCL12, was identified as a candidate interactor. MIF/CXCL4L1 complexation was verified by co-immunoprecipitation, surface plasmon-resonance analysis, and microscale thermophoresis, which also established high-affinity binding (K_D_≍100-150 nM). The binding interface was predicted by peptide array-based mapping and molecular docking. We next determined whether heterocomplex formation modulates inflammatory and atherogenic activities of MIF. MIF-elicited T-cell chemotaxis as assessed in a 3D-matrix-based live cell-imaging set-up was abrogated, when cells were co-incubated with MIF and CXCL4L1. Heterocomplexation also blocked MIF-triggered migration of Egfp^+^ microglia in cortical cultures *in situ*. Of note, CXCL4L1 blocked the binding of Alexa-MIF to a soluble ectodomain mimic of CXCR4 and co-incubation with CXCL4L1 attenuated MIF-triggered dynamic mass redistribution in HEK293-CXCR4 transfectants, indicating that complex formation interferes with MIF/CXCR4 pathways. As MIF and CXCL4L1 are abundant platelet products, we finally tested their role in platelet activation. Multi-photon microscopy, FLIM- FRET, and proximity ligation assay visualized heterocomplexes in platelet aggregates and clinical human thrombus sections. Moreover, heterocomplex formation inhibited MIF- stimulated thrombus formation under flow and skewed the morphology of adhering platelets from a large to a small lamellipodia phenotype. Together, our study establishes a novel molecular interaction, adding to the complexity of the chemokine interactome and chemokine/receptor network. MIF/CXCL4L1, or more generally, ACK/CXC-motif chemokine heterocomplexes may be promising target structures to modulate inflammation and thrombosis.

## Introduction

Chemokines orchestrate immune cell trafficking in health and disease (Charo & Ransohoff, 2006; Noels *et al*, 2019; Weber & Noels, 2011). Chemokine-directed targeting strategies are pursued in acute and chronic inflammatory conditions, autoimmunity, cancer, and athero- sclerosis (Hutchings *et al*, 2017; Noels *et al*., 2019; Zlotnik *et al*, 2011). The chemokine network encompasses 49 classical chemokines (CKs) and 18 classical chemokine receptors (CKRs), which belong to the class of Giα protein-coupled receptors (GPCRs) (Bachelerie *et al*, 2014a; Bachelerie *et al*, 2014b; Murphy *et al*, 2000). Depending on the particular chemokine ligand/receptor pair and various disease and microenvironmental factors, chemokine signaling through CKRs overall capitalizes on the principles of specificity, promiscuity, and biased agonism. Accordingly, multiple chemokines can bind to a certain chemokine receptor and *vice versa*, while ‘biased agonism’ can occur on a ligand, receptor, or tissue basis (Eiger *et al*, 2021; Kleist *et al*, 2016; Steen *et al*, 2014). Fine-tuning of chemokine responses within this network is further expanded by five atypical chemokine receptors (ACKRs) that serve as decoy receptors and promiscuously bind many chemokines to shape their gradients, but also elicit specific signaling responses (Nibbs & Graham, 2013).

Chemokines are well-known to form homodimers, but the discovery of the chemokine interactome additionally suggested a multitude of heteromeric chemokine-chemokine inter- actions even across CC- and CXC-chemokine class borders (Koenen *et al*, 2009; von Hundelshausen *et al*, 2017). CC-type heterodimers between CCL5 and CCL17 or CCL5 and CXCL4 (also termed platelet factor 4, PF4) were found to lead to functional synergism by receptor retention or auxiliary proteoglycan binding and enhancement of chemotactic responses, respectively, while CXC-type heterodimers between CXCL12 and CCL5 or CXCL12 and CXCL4 led to signaling inhibition. This has demonstrated yet another level of complexity within the chemokine network and offers novel intervention strategies in inflammatory and cardiovascular diseases (von Hundelshausen *et al*., 2017).

Moreover, some alarmin-like inflammatory mediators such as human β-defensins (HBDs) and secreted fragments of amino acyl tRNA-synthetases (AARSs), which do not belong to one of the four structural classes of CC-, CXC-, CX_3_C-, or C-chemokines, can bind to chemokine receptors by molecular mimicry and exhibit chemokine-like activities (Rohrl *et al*, 2010; Wakasugi & Schimmel, 1999). These proteins are also referred to as atypical chemokines (ACKs) (Degryse & de Virgilio, 2003; Kapurniotu *et al*, 2019; Oppenheim & Yang, 2005).

Macrophage migration inhibitory factor (MIF) is an evolutionarily conserved pleiotropic inflammatory cytokine (David, 1966; Michelet *et al*, 2019). MIF is an upstream regulator of the host innate immune response and, when dysregulated, is a pivotal mediator of inflammatory diseases, autoimmunity, cancer, and cardiovascular diseases (Calandra & Roger, 2003; Tilstam *et al*, 2017). MIF is a structurally unique cytokine (Sun *et al*, 1996) and, contrary to its eponymous name, has chemokine-like activities and functions as a prototypical ACK (Bernhagen *et al*, 2007; Kapurniotu *et al*., 2019). Accordingly, MIF not only signals through its cognate receptor CD74/invariant chain, but engages in high-affinity interactions with the CXC chemokine receptors CXCR2 and CXCR4 to promote atherogenic monocyte and T-/B-cell recruitment, cancer metastasis, and inflammation (Bernhagen *et al*., 2007; Kapurniotu *et al*., 2019; Klasen *et al*, 2014; Leng *et al*, 2003; Pawig *et al*, 2015; Sinitski *et al*, 2019; Tillmann *et al*, 2013). We elucidated the structural determinants of the binding interface between MIF and its CXC-motif chemokine receptors and found that MIF mimics chemokine receptor binding regions such as the ELR motif and the N-loop (Kraemer *et al*, 2011b; Krammer *et al*, 2021; Lacy *et al*, 2018; Rajasekaran *et al*, 2016; Weber *et al*, 2008). Interestingly, CXCL12/SDF-1α (stromal-derived factor-1α), the cognate ligand of CXCR4, was recently found to bind to the non-chemokine proteins galectin-3 (Eckardt *et al*, 2020) and high-mobility group box-1 (HMGB1) (De Leo *et al*, 2019; Schiraldi *et al*, 2012), but potential interactions between MIF and CXCL12 or CXCL8, the cognate ligand of its chemokine receptors CXCR2, have remained unclear.

Here, we hypothesized that chemokine-chemokine interactions are not only possible between different types of classical chemokines, as demonstrated by chemokine interactome mapping (von Hundelshausen *et al*., 2017), but might extend to ACKs. Choosing MIF as a prototypical ACK, we thus asked whether this mediator would form heterocomplexes with classical chemokines. We tested this hypothesis applying an unbiased chemokine protein array and validated candidate interactors by a battery of biochemical and biophysical methods. We identified the platelet chemokine CXCL4L1 (also termed PF4var1), but not its variant CXCL4, nor the CXCR2 ligand CXCL8 or the CXCR4 ligand CXCL12, as a high affinity interactor of MIF and tested the potential functional role of CXCL4L1/MIF heterocomplex formation for MIF binding to its receptor CXCR4, and in cell systems that are relevant for the inflammatory, atherogenic, and thrombogenic activities of MIF. Finally, we also asked whether such heterocomplexes can be detected in clinical thrombus specimens. Our study extends the chemokine interactome to ACK/CK interactions and demonstrates a functional role for the MIF/CXCL4L1 heterocomplex in disease-relevant activities.

## Materials and Methods

### Proteins and reagents

Biologically active and endotoxin-free recombinant human MIF was prepared as previously described and was obtained at a purity of ∼98% as confirmed by SDS-PAGE analysis in combination with silver staining (Bernhagen *et al*, 1994; Kontos *et al*, 2020). For the preparation of Alexa Fluor-488- and MST-Red-labeled MIF, a 90-95% pure MIF fraction was used. Alexa Fluor-488-labeled MIF was generated using the Microscale Protein Labeling Kit from Invitrogen-Molecular Probes (Karlsruhe, Germany) and MST-Red-MIF was prepared using the Monolith Protein Labeling Kit RED-NHS 2^nd^ Generation from NanoTemper (Munich, Germany), following the manufacturers’ instructions. Biotinylated human MIF was produced using D-biotinoyl- ε -aminocaproic acid-N-hydroxy-succinimide ester (Biotin-7-NHS) with the Biotin Protein Labeling Kit from Roche (Mannheim, Germany). Alternatively, biotin- amidohexanoic acid N-hydroxysuccinimide ester from Sigma-Aldrich (Taufkirchen, Germany) was used.

For the fluorescence polarization assay, a hexahistidine-tagged variant of CXCL4L1 was used. Briefly, the coding sequence of human CXCL4L1 with a methionine-flanked N-terminal His_6_-tag was cloned into the pET21a vector for recombinant bacterial expression using *Xho*I and *Nde*I restriction sites. This construct was then used to transform Rosetta- gami™ 2 (DE3) competent *E. coli* (Novagen, Merck KGaA, Darmstadt, Germany) for subsequent recombinant protein production following induction with 1 mM IPTG (Carl Roth, Karlsruhe, Germany) according to a standard protocol. For purification, bacteria pellets were resuspended in lysis buffer (20 mM Tris-HCl, pH 8.0, 0.5 M NaCl, 5 mM EDTA, 0.1% Triton X-100, with added protease inhibitor tablets according to manufacturer’s instructions) and cells disrupted in a EmlsiFlex-C5 high pressure homogenizer (Avestin Europe GmbH, Mannheim, Germany), the raw extract cleared via centrifugation at 20.000 × g and the resulting pellet, containing recombinant His-CXCL4L1 in inclusion bodies, was washed in lysis buffer with and without detergent. Inclusion bodies were solubilized by gentle shaking overnight in 50 mM Tris-HCl, pH 8.0, 6 M guanidine-HCl, 0.5 M NaCl, 10 mM DTT and His- CXCL4L1 purified from the solubilized pellet via IMAC on a HisTrap HP column on an ÄKTA Pure 25 M FPLC system (Cytiva Europe GmbH, Freiburg, Germany). The obtained protein was subjected to two dialysis steps in refolding buffer (50 mM Tris-HCl, pH 8.0, 0.5 M NaCl, 5 mM methionine, 5 mM cysteine) with and subsequently without 0.9 M guanidine-HCl, followed by a final purification step by size exclusion chromatography in 20 mM sodium phosphate buffer, pH 7.4, using a Superdex 75 10/300 GL column (Cytiva Europe GmbH) on an ÄKTA Pure 25 M FPLC system. A purity degree of 90-95% was verified by SDS-PAGE followed by Coomassie staining and Western Blot according to standard protocols.

Recombinant human peroxiredoxins 1 and 6 (PRX1, PRX6) were purchased from Abcam (Abcam PLC, Cambridge, UK), while recombinant human β-defensin-1 and 2 (HBD-1, HBD-2) were obtained from ProSpec (ProSpec-Tany TechnoGene Ltd., Ness Ziona, Israel). Recombinant human HMGB1 was purchased from Novus (Novus Biologicals Europe, Abingdon, UK). Recombinant human CXCL4L1 (PF4var1) as well as the CXCL4 (PF4) were purchased from ChromaTec (Greifswald, Germany). The other recombinant human chemokines were obtained from Peprotech (Hamburg, Germany). All other reagents and chemicals were purchased from Merck KGaA (Darmstadt, Germany), Carl Roth GmbH (Karslruhe, Germany), or Sigma-Aldrich and were of the highest purity degree available.

### Cell culture and cultivation of mammalian cell lines

Jurkat T cells were cultured in RPMI1640 medium (Gibco) supplemented with 10% fetal calf serum (FCS), 1% penicillin/streptomycin, and 1x non-essential amino acids (NEAAs, Gibco). The human monocytic cell line MonoMac6 (Ziegler-Heitbrock *et al*, 1988) was cultured in RPMI1640 medium + GlutaMAX (1x), supplemented with 1x NEAAs, 10% FCS, and 1% penicillin/streptomycin. HEK293 cells stably transfected with human CXCR4 (HEK293- CXCR4) were used at passage 5 and were cultivated in DMEM medium (Gibco), supplemented with 10% FCS and 1% penicillin/streptomycin (Gibco), and used for the experiment between passage 6 and 8.

Unless stated otherwise, cells were cultivated in a temperature- and humidity-controlled incubator at a temperature of 37°C and 5% CO_2_. FCS from an EU-approved origin was obtained from Invitrogen-Thermo Fisher Scientific and heat-inactivated prior to usage. Other cell culture reagents, media and supplements were bought from Invitrogen-Thermo Fisher Scientific, unless stated otherwise. Cell lines were originally obtained from the German Society for Microorganisms and Cell Cultures (DSMZ, Braunschweig, Germany) or from the American Type Culture Collections (ATCC).

### Isolation of primary human CD4^+^ T cells

Primary human CD4-positive T cells were isolated from enriched peripheral blood mononuclear cell (PBMC) fractions using the human CD4^+^ T cell isolation kit from Miltenyi Biotec (Bergisch Gladbach, Germany) according to the manufacturer’s instructions. Cells were cultivated in RPMI1640 medium, supplemented with 10% FCS, 1% penicillin-/streptomycin, and 1x NEAAs in a cell culture incubator at 37°C and 5% CO_2_ and used for functional assays on the next day. PBMC fractions were obtained by apheresis from conical chambers of a Leucoreduction System Chamber sourced from anonymous platelet donations at the Department of Transfusion Medicine, Cell Therapeutics and Hemostaseology of LMU University Hospital. Studies abide by the Declaration of Helsinki principles and were approved by ethics approval 18-104 of the Ethics Committee of LMU Munich, which encompasses the use of anonymized tissue and blood specimens for research purposes.

### Isolation of human platelets

#### For immunofluorescent stainings

Human platelets were isolated from blood, freshly drawn from healthy donors, using a syringe containing 1/10 volume of CTAD–buffer (0.105 M tri-sodium citrate, 10 mM theophylline, 3.7 mM adenosine, 0.198 mM dipyridamole) (Polack *et al*, 2001). To prevent platelet activation, the blood was supplemented with prostaglandine E1 (Merck KGaA), Apyrase (New England Biolabs GmbH, Frankfurt am Main, Germany), and EGTA (Sigma-Aldrich). Briefly, platelets were isolated by sequential centrifugation steps, performed at room temperature (RT) with reduced brake settings. Platelet-rich plasma (PRP) was separated from whole blood by centrifugation for 5 min at 300 × g, diluted with an equal volume of phosphate-buffered saline (PBS), pH 7.4, and centrifuged again for 10 min at 200 × g to remove remaining leukocytes. Finally, platelets were sedimented by centrifugation for 10 min at 400 × g.

#### For functional studies

Washed human platelets were isolated as previously described (Borst *et al*, 2012) and subsequently used for functional flow chamber or platelet spreading assays.

### Mice and preparation and cultivation of primary mixed cortical cultures for the microglia motility assay

CX3CR1^GFP/+^ mice, which were originally obtained from the Jackson Laboratories (strain 005582; (Niess *et al*, 2005)), were established on a pure C57BL/6 background and housed under standardized light-dark cycles in a temperature-controlled air-conditioned environment under specific pathogen-free conditions at the Center for Stroke and Dementia Research (CSD), Munich, Germany, with free access to food and water. Animals were sacrificed under anaesthesia with a mixture of midazolam (5 mg/mL), medetomidine and fentanyl (MMF). Mouse maintenance and experiments were reviewed and overseen by the institutional animal use and care committee of the local authorities (Regierung von Oberbayern, ROB, Germany) and performed in accordance with the procedures provided by the animal protection representative of CSD.

Primary mixed cortical cultures containing CX3CR1^GFP/+^ microglia were prepared in 96-well imaging plates based on a previously established protocol (Gokce & Sudhof, 2013) from the cortices of 5 newborn pups of the CX3CR1^GFP/+^ mouse line (postnatal day 0) in plating medium, consisting of modified Minimum Essential Medium (MEM without glutamine and phenol red) (Gibco), supplemented with 0.5% glucose, 0.02% sodium bicarbonate, 1x ITS-supplement (Sigma-Aldrich), 2 mM L-glutamine (Gibco), 1% penicillin/streptomycin, and 10% FCS. Cultures were incubated in a humidified atmosphere at 37 °C and 5% CO_2_ for 10 d. One day after plating, 80% of the plating medium was replaced with growth medium, prepared from MEM (without glutamine and phenol red) supplemented with 0.5% glucose, 0.02% sodium bicarbonate, 5% FCS, 0.5 mM L-glutamine, and serum-free B-27™ supplement (Gibco). On the fourth day after dissection, 50% of the medium was replaced with growth medium additionally supplemented with 4 μM cytosine-1-β-D-arabinofuranoside (Sigma-Aldrich).

### Chemokine protein array

Human chemokines and selected atypical chemokines were spotted on a nitrocellulose membrane at 100 ng per spot and left to dry at RT. Membranes were blocked with 1x ROTI®Block (Carl Roth) for 2 h at RT and then probed overnight with biotinylated human MIF (biotin-MIF) at a concentration of 1 µg/mL in either 10 mM Tris-HCl pH, 8.0 or 10 mM MES, pH 6.0. Subsequently, membranes were washed three times with 0.01% Tween®20 in water and developed with horseradish-peroxidase (HRP)-conjugated streptavidin (Bio-Techne GmbH, Wiesbaden-Nordenstadt, Germany), diluted 1:200 in 1x ROTI®Block, for 2 h. After another washing step, bound biotin-MIF was revealed via chemiluminescence using SuperSignal™ West Pico Chemiluminescent Substrate (Thermo Fisher Scientific) on a LAS- 3000 Imaging System (Fuji Photo Film Co., LTD., Japan).

### Pull-down of CXCL4L1 from cell lysates

MonoMac-6 cells were first washed with PBS and then lysed on ice for 30 min with immunoprecipitation (IP) lysis buffer (1x cell lysis buffer, Cell Signaling, cat#9803), 100 mM PMSF, and 1x protease and phosphatase inhibitors (ThermoFisher). The purified cell lysates were then incubated with pre-washed streptavidin-conjugated paramagnetic beads (DYNAL™ Dynabeads™ M-280 Streptavidin; Invitrogen, cat#11205D) for 2 h at 4 °C (preclearing step). After centrifugation, the supernatant was incubated with biotinylated human MIF by gentle, constant shaking on a rotary shaker overnight at 4 °C. To capture MIF/CXCL4L1 complexes, prewashed streptavidin-conjugated beads were added to the precleared lysates, and the mixture was incubated for 2 h at 4 °C on a rotary shaker. Beads were separated from the lysate using a magnetic stand (Dynal™ MCP-S) and washed three times with lysis buffer. The supernatant was removed and the beads were resuspended in LDS sample buffer (Invitrogen) and boiled at 95 °C for 15 min. Samples were subjected to SDS-PAGE and analyzed by Western blotting. For this purpose, equal amounts of protein were loaded onto 11% SDS-polyacrylamide gels (NuPAGE, Thermofisher) and transferred to polyvinylidene difluoride (PVDF) membranes (Carl Roth, Karlsruhe, Germany). Membranes were blocked in PBS-Tween-20 containing 5% BSA for 1 h and incubated overnight at 4 °C with rabbit polyclonal anti-MIF antibody Ka565 (Bernhagen *et al*., 2007) or rabbit polyclonal anti-PF4V1 IgG PA5-21944 (Invitrogen) diluted in blocking buffer. Proteins were revealed using anti-rabbit HRP as a secondary antibody. Signals were detected by chemilumi- nescence on an Odyssey® Fc Imager (LI-COR Biosciences GmbH, Bad Homburg, Germany) using SuperSignal™ West Dura ECL substrate from ThermoFisher Scientific and specific primary antibodies as indicated.

### CelluSpot peptide array

The CelluSpot peptide array method has been described previously (Lacy *et al*., 2018). Briefly, 15-meric peptides, positionally frame-shifted by three residues and spanning the entire sequence of CXCL4 and CXCL4L1, were synthesized on modified cellulose disks (Intavis MultiPep RSi/CelluSpot Array, Cologne, Germany). Peptides were then further processed by dissolving the cellulose, and spotted on coated glass slides using a slide spotting robot from Intavis. Slides were incubated in blocking buffer (50 mM Tris-buffered saline, pH 7.4, 1% BSA, 0.1% Tween® 20), washed (50 mM Tris-buffered saline, pH 7.4, 0.1% Tween® 20) and probed with biotinylated human MIF (3 µM in blocking buffer). After washing, slides were developed with a dilution of streptavidin-conjugated horseradish peroxidase (Roche) in blocking buffer. Bound MIF was revealed by chemiluminescence on an Odyssey® Fc imager using the SuperSignal™ West Dura ECL substrate

### Microscale thermophoresis (MST)

Protein-protein interactions were analyzed via microscale thermophoresis on a Monolith NT.115 instrument equipped with green/red filters (NanoTemper Technologies, Munich, Germany). Measurements were performed at 25°C at both 40% and 80% MST power. LED excitation power was adjusted to 90 or 95% in order to obtain an initial fluorescence count of 700 to 800. MST traces were recorded for 40 s (-5 s to +35 s), according to default settings with the sample being heated from 0 to 30 s. All measurements were performed in assay buffer (10 mM Tris-HCl, pH 8.0, 0.01% BSA). MST-Red-MIF was used at a fixed concentration, mixed 1:1 with serial dilutions of either CXCL4 (Peprotech, Hamburg, Germany) or CXCL4L1 (ChromaTec, Greifswald, Germany) (final MIF concentrations: 456 nM or 312 nM, respectively). Prior to measurement, the prepared samples were incubated for at least 30 min on ice. MST traces of multiple experiments were analyzed according to the K_D_ model using the default T-jump settings, focusing on the temperature related intensity change (TRIC) of the fluorescent label (“cold region” from -1 to 0 s, “hot region” from 0.5 t 1.5 s) using the MO.AffinityAnalysis V2.3 software (NanoTemper Technologies). Curve fitting for data presentation was performed by GraphPad Prism Version 6.07 (‘one site – total binding’).

### Analysis of protein-protein interactions by surface plasmon resonance (SPR)

Surface plasmon resonance measurements were performed using a Biacore X100 instrument (GE Healthcare Europe GmbH) and neutravidin-modified C1 sensor chips. Biotin-MIF was immobilized on flow cells to 1064.8 RU. CXCL4 (Peprotech, Hamburg, Germany) and CXCL4L1 (ChromaTec, Greifswald, Germany), used at concentrations in the range of 0.125 to 20 µg/mL in running buffer (HBS-EP+ Buffer: 0.01 M HEPES, 0.15 M NaCl, 0.003 M EDTA and 0.05% v/v surfactant P20) were injected at a flow rate of 60 µL/min. The complex was allowed to associate and dissociate for 90 s and 240 s, respectively. Surfaces were regenerated with 2 pulses (60 s) of 30 mM NaOH and 2 M NaCl. Responses from analyte injections were fitted to a 1:1 Langmuir interaction profile using Biacore X100 evaluation 2.0.1 Plus package software.

### Transwell migration assay

Transwell migration experiments to study the influence of CXCL4L1 on MIF-mediated chemotaxis responses were performed with Jurkat T cells. Briefly, Jurkat cells were diluted in RPMI1640 medium at a density of 1 × 10^7^ cells/mL. Cells were placed in the upper chamber of a 24-well Transwell insert with 5μm pore size (Corning, Kaiserslautern, Germany). 16 nM MIF, either alone or pre-incubated (30 min on ice to allow for complex formation) with 32 nM of CXCL4L1, as well as 32 nM CXCL4L1 alone were added to the lower chamber as a chemoattractant. After a 12 h migration interval at 37 °C and 5% CO_2_, migrated cells were recovered from the lower chamber and counted via flow cytometry by using CountBright™ absolute counting beads (Molecular Probes-Invitrogen). In a similar experimental setup, the influence of CXCL4 on MIF-mediated chemotaxis was tested as well. MIF was used at a concentration of 16 nM and CXCL4 (ChromaTec, Greifswald, Germany) at 32 nM.

### 3D migration of human CD4^+^ T cells

The migratory behavior of primary human T cells was assessed by three-dimensional (3D) migration methodology using time-lapse microscopy and single cell tracking using the 3D chemotaxis µ-Slide system from Ibidi GmbH (Munich, Germany). The method was performed following a slight modification of the established Ibidi dendritic cell protocol for human monocytes, as described previously (Kontos *et al*., 2020). Briefly, isolated CD4^+^ human T cells (3.5 × 10^6^) were seeded in a rat tail collagen type-I gel (Ibidi, Munich, Germany) in DMEM and subjected to a gradient of human MIF, CXCL4L1, or a pre-incubated combination of both. Cell motility was monitored performing time-lapse imaging every 0.5 or 2 min at 37°C for a period of 120 min to cover either a short or extended migration period, using a Leica DMi8 inverted microscope (Leica Microsystems, Wetzlar, Germany) and Leica live cell-imaging software (LAS X version 3.7.4). Images were imported as stacks to ImageJ version 1.51n and analyzed with the manual tracking and Chemotaxis and Migration tool (Ibidi GmbH) plugin for ImageJ.

### Motility measurement of primary murine microglia

The motility of mouse microglia was determined using mixed cortical cultures, established and cultivated as stated above. A day prior to imaging, the medium was changed to Hibernate A medium (Gibco) in order to maintain a physiological pH value during imaging. Prior to imaging, different wells of cells from each individual pup were treated with either 8 nM MIF, 1.6 nM CXCL4L1, or both (pre-incubated for 30 min on ice to allow for complex formation). A control group was treated with 20 mM sodium phosphate buffer, pH 7.4. Cell motility was monitored by time-lapse imaging for 15 h at 37 °C with recordings every 5 min, using a Leica DMi8 inverted Life Cell Imaging System using the FITC channel for visualizing the GFP-positive cells. Images were imported as stacks to ImageJ software version 1.51n and analyzed with the manual tracking and Chemotaxis and Migration tool Plugin for ImageJ from Ibidi. In order to quantify microglial motility from the time-lapse videos, 20-25 GFP-positive microglia per treatment group were randomly selected and manually tracked throughout all frames. Cells that died or moved out of the frame were excluded from the analysis. Accumulated distance of each tracked microglia was calculated with Chemotaxis and Migration Tool (Ibidi).

### Fluorescence polarization spectroscopy

Fluorescence polarization was measured using a JASCO FP-6500 fluorescence spectrophotometer equipped with FDP-223 and FDP-243 manual polarizers (JASCO Deutschland GmbH, Pfungstadt, Germany). Preparation of stock solutions, measurements and analysis were performed essentially following a previously published protocol (Kontos *et al*., 2020). For binding/inhibition experiments, mixtures of Alexa 488-labeled MIF (10 nM) in the absence/presence of CXCL4L1 (1.6 µM) (or 20 mM sodium phosphate buffer, pH 7.2), and non-labeled msR4M-L1 peptide (concentration between 1 nM to 10 µM) were prepared in 10 mM sodium phosphate, pH 7.2, containing 2% hexafluoro-isopropanol (HFIP). Where CXCL4L1 was added as a putative inhibitor, Alexa488-MIF and CXCL4L1 were mixed and incubated for 30 min prior to measurements. Bandwidth for excitation and emission was set at 5 nm and time response at 0.5 s. The excitation wavelength was 492 nm and emission was recorded at 519 nm. Measurements were taken at RT within 2 to 3 min upon preparation of the solutions. Polarization P was calculated according to the equation P = (I_ − G·I_⊥_) / (I_ + G·I_⊥_), with I_ as the intensity of emitted light polarized parallel to the excitation light and I_⊥_ as the intensity of emitted light polarized perpendicular to the excitation light. The G factor was calculated based on the instrumental documentation (Moerke, 2009). Apparent K_D_ values were calculated assuming a 1:1 binding model (Yan *et al*, 2006), using sigmoidal curve fitting with OriginPro 2016 (OriginLab Corporation, Northampton, MA, USA).

### Label-free dynamic mass redistribution (DMR) assay

Analysis of dynamic mass redistribution of adherent cells was performed on an EnSpire Multimode plate reader equipped with an Epic® label-free measurement module (PerkinElmer Inc., Waltham MA, USA) according to the manufacturer’s instructions for cell-based label-free DMR measurements (Schroder *et al*, 2010). The assay protocol was adapted according to a previous publication to be performed in EnSpire label-free 96-well fibronectin-coated cell assay microplates (Corning GmbH, Amsterdam, The Netherlands) with HEK293 cells stably expressing human CXCR4 (Krammer *et al*., 2021). Briefly, 40.000 cells were seeded into each well and cultivated overnight (37°C, 5% CO_2_) to achieve a confluency of >70%. Prior to the assay, the medium was exchanged with DMR assay buffer (20 mM HEPES and 1% DMSO in HBSS, pH 7.4) and the assay plate left for 6 h to equilibrate to ambient temperature. Baseline measurements for each well were recorded for 10 min every 30 s prior to treatment of the cells with chemokines, inhibitors, or the corresponding buffers as control. Treatments were applied to each well as a 5x concentrated stock, prepared in assay buffer. For treatment with MIF/CXCL4L1 complexes, both proteins were mixed in assay buffer and incubated for 5 h at RT. Directly after addition of the stimuli, the DMR response was recorded for the indicated duration. The DMR response resembles the wavelength shift of the light reflected from the sensor integrated in the assay microplates and serves as a cumulative cellular response signal. Measurements were performed on two replicates per treatment and results are presented as their mean value.

### Staining of human thrombus specimens

Human thrombus tissue specimens, obtained as disposable material from vascular surgery procedures (ethics allowance LMU Munich 18-104 and TUM-MRI project # 2799/10), were embedded in Tissue Tek O.C.T. Compound (Sakura Finetek Germany GmbH, Staufen, Germany), frozen, and cut into 5 µm sections using a CM 1950 Cryostat (Leica Biosystems). The cryosections were transferred to microscopy slides and stored at -80°C until use.

#### Hematoxylin & eosin (HE) staining of thrombus sections

Cryosections of human thrombus tissue were stained with Mayer’s hematoxylin and eosin (HE) according to standard protocols. Briefly, after thawing and brief air drying and rehydration, sections were incubated in Mayer’s hematoxylin solution (Sigma-Aldrich) for 15 min. After thorough rinsing of the samples, 0.5% eosin Y solution (Sigma-Aldrich) was applied as a counterstain for 30 s. After dehydration of the tissue in 95 % ethanol, 100 % ethanol (Merck KGaA) and xylene (VWR International GmbH), the sample was covered with resinous mounting medium (Eukitt, Sigma-Aldrich), covered with a glass coverslip and examined by light microscopy (Leica Dmi8 inverted microscope using a DMC2900 digital camera (Leica Microsystems; 10x objective).

### Staining of platelets

Freshly isolated platelets were fixed with 4% paraformaldehyde (PFA) in PBS (Morphisto GmbH, Frankfurt a. M., Germany) for 10 min and subsequently permeabilized using 1x Perm buffer (Invitrogen) for 15 min. After washing, platelets were blocked in ROTI®Block (Carl Roth) for 1 h. Immunofluorescent staining was performed as described above for the human thrombus specimens, except that the anti-human MIF antibody was used at a dilution of 1:20 and the anti-human CXCL4L1 antibody at a dilution of 1:50. Stained platelets were then washed in blocking buffer and mounted on poly-L-ornithine-coated glass slides using ProLong^TM^ Glass Antifade mountant (Invitrogen), covered with coverslips and stored at 4 °C until imaging by multiphoton microscopy.

### Multiphoton laser-scanning microscopy (MPM) and FLIM-FRET

Imaging was conducted using a multispectral TCS SP8 DIVE FALCON LIGHTNING microscope (Leica, Germany) equipped with filter-free 4TUNE NDD detection module, an extended IR spectrum tunable laser (New InSight® X3™, Spectra-Physics) (680-1300 nm) and fixed IR laser (1045 nm), advanced Vario Beam Expander (VBE), Ultra-high-speed resonance scanner (8kHz), HC PL IRAPO 25x/1.0 WATER objective, and FLIM-FRET modality. Images were collected in a sequential scanning mode using hybrid diode detectors Reflected Light Hybrid Detectors (HyD-RLD) (Alexa Fluor-488: excitation 965 nm / emission 479-568 nm; Cy3: excitation 1095 nm / emission 538-650 nm) and were handled using the LAS-X software package. Deconvolution microscopy was performed using the Leica LIGHTNING (adaptive deconvolution) application.

For fluorescence lifetime imaging (FLIM) and FLIM-FRET measurements, up to 1000 photons per pixel were captured in a time-correlated single photon counting (TCSPC) mode. Fluorescence lifetime decay data were fitted using Leica FALCON (FAstLifetime CONtrast) software. The fitting was assessed by randomly distributed residuals and by low Chi-square (χ^2^) values. The number of components used for the fittings was manually fixed to a value (n=2-3) to minimize χ^2^values. The fluorescence lifetime of the donor was acquired similarly in the absence of the acceptor.

### Proximity ligation assay (PLA)

For detection of protein complexes by proximity ligation assay (PLA), the Duolink^TM^ InSitu Orange Starter Kit Mouse/Rabbit (DUO92102) from Sigma Aldrich was used. Following scouting experiments to establish the PLA methodology in thrombus material, cryosections of the thrombi were prepared by treatment with cold acetone for 6 min at 4 °C and for 30 min at RT. Samples were rehydrated in PBS for 10 min and hydrophobic barriers were applied to the microscopy slide using an ImmoEdgeTM Pen (Vector Laboratories Inc. Burlingame, USA).

For PLA detection, the Duolink® PLA Fluorescence protocol provided by the manufacturer was essentially followed, using primary antibodies against human MIF (mouse anti-MIF D2, sc-271631, Santa Cruz Biotechnology Inc., Dallas, USA; 1:20) and against human CXCL4L1 (rabbit anti-CXCL4L1, PA5-21944, Invitrogen; 1:50). Samples were then prepared for microscopy using Duolink® mounting medium with DAPI, and coverslips sealed with commercially available nail polish and stored at 4 °C until imaging by confocal microscopy using a Zeiss LSM880 AiryScan microscope was performed.

### Flow chamber assay with platelets

Chemokines were diluted in calcium-free PBS, pH 7.4, at their final concentrations (MIF: 16 nM; CXCL4L1: 32 nM) and allocated into separate reaction tubes. 200 µL of each solution were distributed onto separate collagen-coated cover slips (100 µg/mL) and incubated for 2 h. Cover slips were blocked with PBS, pH 7.4, containing 1% BSA for 1 h. Next, human whole-blood was diluted at a 5:1 ratio with PBS, pH 7.4, containing calcium. Before perfusion, the blood was incubated with fluorochrome 3,3 -dihexyloxacarbocyanine iodide (DiOC_6_, 1 mM; Sigma Aldrich) for 10 min at RT. Thereafter, the blood was allocated into 1 mL syringes and perfused over the different cover slips, through a transparent flow chamber with high shear rate (1000 s^−1^) for 5 min. Per run, one 2-minute video clip was recorded (200 ms/frame, Nikon Eclipse Ti2-A, 20x objective). Afterwards, the chamber was rinsed and pictures were taken of five representative areas using the same objective. The covered area was analyzed using the NIS-Elements AR software (Nikon) and the mean percentage of the covered area, the mean thrombus area as well as the mean thrombus count were determined.

### Platelet spreading analysis

Fibrinogen-coated (100 µg/mL, Sigma Aldrich) coverslips were preincubated with MIF (16 nM), CXCL4L1 (32 nM), or MIF (16 nM) and CXCL4L1 (32 nM) together, for 2 h. Afterwards, isolated human platelets were diluted in Tyrodes buffer (pH 7.4) to match a concentration of 15.000 cells/µL. Platelets were supplemented with 1 mM CaCl_2_, activated with 1 µg/mL CRP-XL (CambCol, Cambridge, UK), and incubated on the previously prepared fibrinogen-coated coverslips for 30 or 60 min at RT. Thereafter, platelets were fixed with 4% formaldehyde (Sigma Aldrich) for 10 min, and washed three times with PBS, pH 7.4. The coverslips were mounted onto slides and five images from randomly selected areas were taken using a Nikon Eclipse Ti2-A microscope with a 100x DIC objective. Subsequently a quarter of each image with at least 20 cells was analyzed.

### Protein structure visualization

Three-dimensional structures as well as the surface charge distribution of human MIF, CXCL4 and CXCL4L1 were visualized using the PyMOL Molecular Graphics System software, version 1.8.2.2 (Schrödinger, LLC). The structures represent the Protein Data Bank (PDB) files for MIF (PDB ID: 3DJH), CXCL4 (PDB ID: 1F9Q), and CXCL4L1 (PDB ID: 4HSV), orour molecular docking results.

### Protein-protein docking

To simulate the interaction of monomeric MIF with CXCL4 and CXCL4L1 in their monomeric forms, rigid protein-protein docking, followed by clustering of the 1000 lowest energy structures and removal of steric clashes was performed using the ClusPro 2.0 webserver, with single chains of MIF and CXCL4L1 defined as ‘receptor’ and ‘ligand’, respectively (Kozakov *et al*, 2017; Vajda *et al*, 2017).

### Statistical analysis

Statistical analysis was performed using GraphPad Prism Version 6.07 software. Unless stated otherwise, data are represented as means ± standard deviation (SD). After testing for normality, data were analyzed either by two-tailed Student’s T-test, Mann-Whitney U test, or Kruskal-Wallis test as appropriate. Differences with *P*<0.05 were considered to be statistically significant.

## Results

### High affinity binding between the atypical chemokine MIF and the platelet CXC chemokine CXCL4L1

To begin to test the hypothesis that chemokine-chemokine interactions may extend to ACKs and that MIF may form heterocomplexes with classical chemokines, we applied unbiased chemokine protein array technology (*Figure 1A-B*), as previously successfully used to map formation of heterocomplexes between different classical chemokines (von Hundelshausen *et al*., 2017). In addition to 47 human chemokines covering all four sub-classes (CXC-, CC-, CX3C- and C-type CKs) we also included structurally related and positively charged protein mediators including ACKs/DAMPs such as HMGB1, HBDs, and peroxiredoxins (Prxs) (Shichita *et al*, 2012; He *et al*, 2019), as well as MIF itself and the MIF homolog D-dopachrome tautomerase (D-DT)/MIF-2 as spotted proteins in the protein array. Probing of the array with biotin-conjugated MIF and streptavidin-POD (StrAv-POD) revealed high-intensity spots indicative of a tight interaction of MIF with CXCL4L1 and Prx1 (*Figure 1B-C*). Weaker spots were detected for CCL28, CXCL9, Prx6, and MIF itself. No spot intensity whatsoever was observed for any of the other immobilized proteins, indicating that none of the other 44 chemokines interacts with MIF. This also included CXCL8 and CXCL12, which share their receptors CXCR2 and CXCR4, respectively, with MIF (Bernhagen *et al*., 2007).

**Figure 1:**
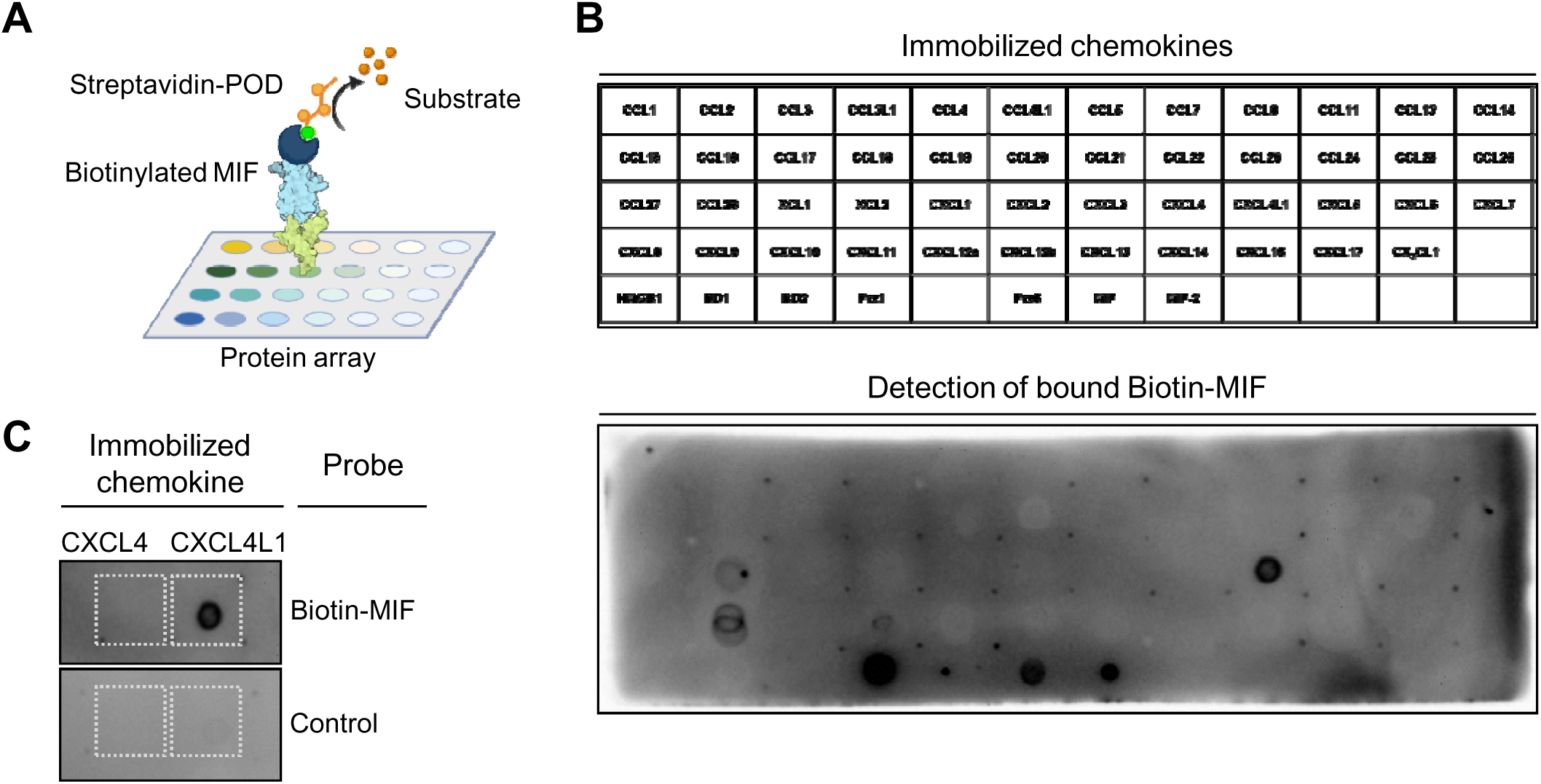
Unbiased chemokine protein array identifies CXCL4L1, but not CXCL4, as a novel interaction candidate of MIF. (**A**) Schematic illustrating binding of biotinylated MIF to the chemokine protein array. (**B**) Layout of the immobilized chemokines, atypical chemokines and alarmins (*top*) and membrane of chemokine solid phase assay performed at pH 8.0, developed against bound biotin-MIF (*bottom*). (**C**) Close-up of the membrane with a focus on CXCL4 and CXCL4L1 with the corresponding negative control membrane, incubated without biotin-MIF.

Similarly, no binding signal of biotin-MIF was detected with HMGB1, a DAMP which has been demonstrated to form heterodimers with CXCL12 and for which a functional interaction with MIF has been suggested (Ma *et al*, 2017; Schiraldi *et al*., 2012), nor for the human β-defensins HBD1 or HBD2 (*Figure 1B-C*). Importantly, when testing a control chemokine array developed with StrAv-POD without a biotin MIF incubation step, only one signal was not fully specific. This was the signal for Prx1 so that its interpretation was not possible (*Supplementary Figure 1A*). Biotin-MIF also bound to MIF itself, but not to MIF-2 (*Figure 1B-C*). As MIF is known to form homo-oligomers (Sun *et al*., 1996) and has been reported to form higher-order hexameric complexes (Bai *et al*, 2012), this result further verified the validity of the chemokine array approach for MIF.

A striking observation was that biotin-MIF specifically interacted with the immobilized platelet chemokine CXCL4L1, but not with CXCL4 (*Figure 1C* and *Supplementary Figure 1B*). CXCL4 and CXCL4L1 are highly homologous chemokines, their sequences only differ by three amino acids, and CXCL4L1 has also been suggested to be a decoy chemokine paralog of CXCL4. Given this remarkable specificity of the interaction with MIF and that the spot corresponding to biotin-MIF and CXCL4L1 was the strongest interaction detected on the array, we focused on CXCL4L1 as a novel candidate interactor of MIF.

We first verified the interaction by co-immunoprecipitation using whole cell lysates of MonoMac6 cells, which we found to express substantial amounts of CXCL4L1. Semi- endogenous pulldown of proteins from MonoMac6 lysates by biotin-MIF and StrAv magnetic beads and Western blot using an anti-human CXCL4L1 antibody revealed a specific band for CXCL4L1, which was absent when the pulldown was performed without biotin-MIF preincubation (*Figure 2A*). Pulldown specificity was further confirmed by Western blot against MIF. We next applied surface plasmon resonance (‘Biacore’) methodology, which was previously successfully used to characterize interactions within the classical chemokine interactome (von Hundelshausen *et al*., 2017). To study the MIF/CXCL4L1 interaction, MIF chips were exposed to increasing concentrations of CXCL4L1 in the soluble phase. The obtained surface plasmon resonance response curves indicated that MIF specifically binds to CXCL4L1 (*Figure 2B-C*). Quantitative analysis determined a K_D_ value of 116 ± 16 nM (mean ± SD) indicating high-affinity binding between MIF and CXCL4L1. By contrast, no appreciable signal was detectable for the incubation with increasing concentrations of CXCL4 and no K_D_ could be derived, verifying the specificity of the MIF/CXCL4L1 interaction in this set-up. To further confirm the MIF/CXCL4L1 interaction, we next applied microscale thermophoresis (MST), which relied on the interaction between MST-Red-labeled MIF and its binding partner, with both partners in the soluble phase. This methodology was recently established for MIF (Kontos *et al*., 2020). MST titrations of MST-Red-MIF with increasing concentrations of CXCL4L1 revealed a typical sigmoidal binding curve with a derived binding constant (K_D_ = 159.8 ± 16.8 nM) that was similar to that obtained by surface plasmon resonance (*Figure 2D-E*). In contrast, binding was much weaker when CXCL4 was titrated and accordingly a low affinity K_D_ in the micromolar range was determined (K_D_ = 2.0 ± 0.8 µM).

**Figure 2:**
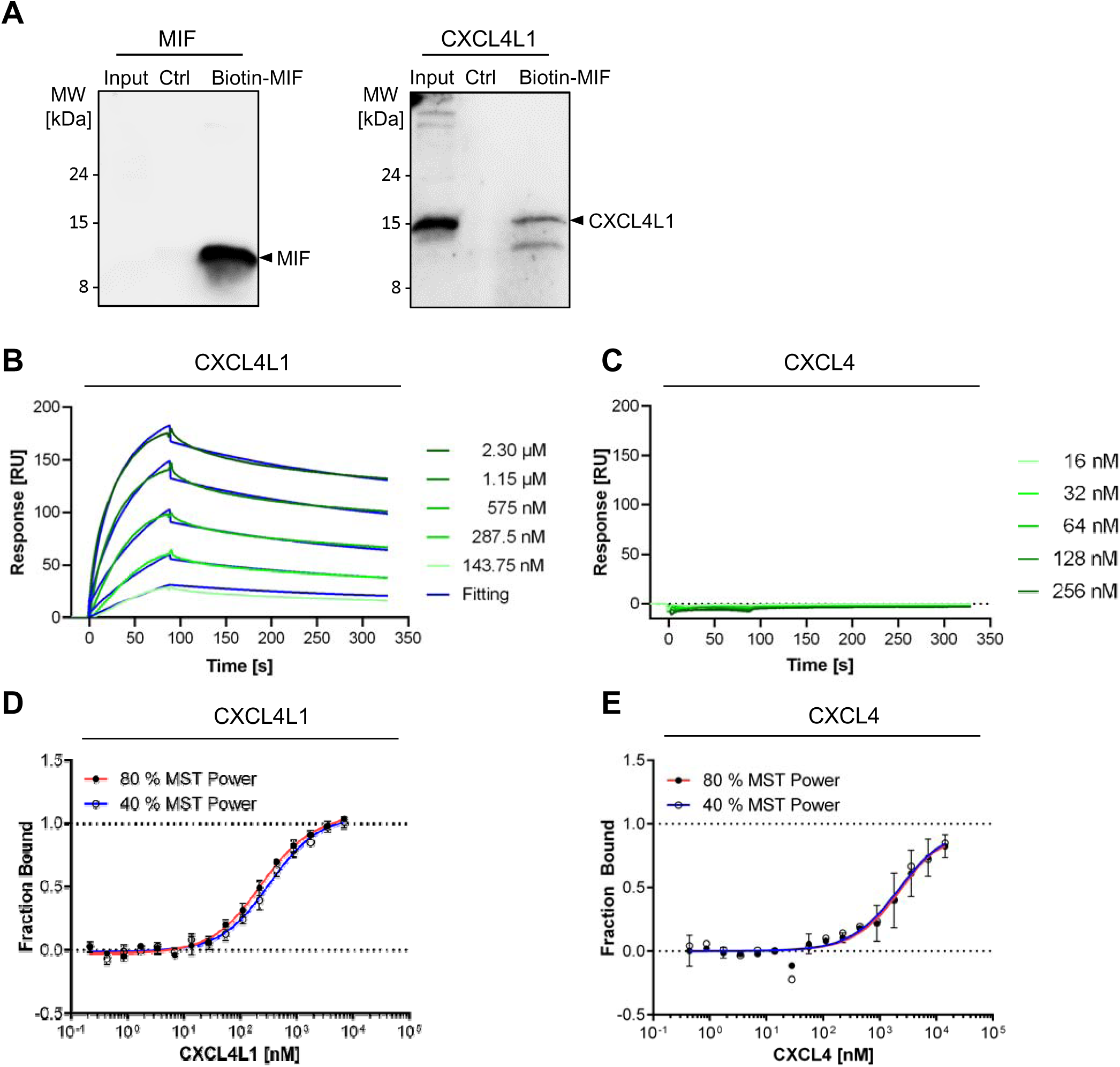
Validation of MIF/CXCL4L1 complex formation by a variety of protein-protein interaction assays and verification of the specificity of MIF complexation with CXCL4L1 over CXCL4. (**A**) Semi-endogenous pull-down assay, in which endogenous CXCL4L1 from MonoMac6 lysates was captured with recombinant biotinylated MIF and pulled down by streptavidin-coated paramagnetic beads. Blots, developed against MIF (*left*) and CXCL4L1 (*right*), show representative results of three independent experiments. Input corresponds to 5% cell lysate without pull-down and control (Ctrl) refers to pull-downs performed in the absence of biotin-MIF. Molecular weight markers were lelectrophoresed in the same gel and relevant marker sizes are indicated. (**B**) Interrogation of MIF/CXCL4L1 complex formation by surface plasmon resonance (SPR) spectroscopy using chip-immobilized biotin-MIF titred against increasing concentrations of CXCL4L1. Measurements indicate an interaction between MIF and CXCL4L1 with an estimated K_D_ of 116 ± 16 nM. The SPR response signal is given in relative units (RU). (**C**) Same as (**B**), except that titration was performed with CXCL4. Corresponding SPR spectroscopy data for MIF and CXCL4. No detectable binding signal was obtained and no K_D_ could be derived. (**D**) Interrogation of MIF/CXCL4L1 complex formation by microscale thermophoresis (MST) utilizing fluorescently labeled MIF and CXCL4L1 in solution. MST analysis revealed a K_D_ of 159.8 ± 16.8 nM for the interaction of MIF and CXCL4L1. (**E**) Same as (**D**), except that CXCL4 was tested. The derived apparent K_D_ of 2.0 ± 0.8 µM was ten-fold higher compared to MIF/CXCL4L1.

Heterodimer formation between classical chemokines relies on CC-type or CXC-type interactions. To determine which residues in CXCL4L1 are critical for the interaction with MIF, we employed peptide array technology. A set of 15-meric peptides derived from the CXCL4L1 sequence, positionally frame-shifted by three amino acids to cover the entire sequence of the processed chemokine, were synthesized and immobilized on glass slides and arrays, and probed with biotin-MIF. The most pronounced binding signal was observed for peptides representing the sequence region, which corresponds to the β2-strand motif IKAGPHCPTAQLIAT of CXCL4L1 (*Supplementary Figure 2A-B*). A second peak encompasses the N-terminal sequence QCLCVKTTSQVRPRH. The difference in the 3D structures of CXCL4 and CXCL4L1 is characterized by a significant conformational rearrangement of the α-helix (Kuo *et al*, 2013), although the sequence of CXCL4 differs from that of CXCL4L1 in only three α-helical residues (L58P, K66E, L67H with the conformational difference being mainly governed by the L67H exchange). In this respect, CXCL4 showed an essentially identical peptide binding profile as that of CXCL4L1 at the N-terminus as expected, but a slightly different pattern at the β2 strand region GPHCPTAQLIATLKN, that is packed onto the C-terminal α-helix (*Supplementary Figure 2A-B*). Peptide array-based mapping of the CXCL4L1 residues involved in MIF binding was confirmed by molecular docking simulations. Docking applying the ClusPro software predicted that the β-sheet region including the IKAGPHCPTAQLIAT motif is located near the MIF contact site, facing the 4- stranded β-sheet of a single MIF monomer chain. This interaction could be promoted by an energetically favorable complementary electrostatic interaction between the two surfaces (*Supplementary Figure 2C*).

Together, the co-immunoprecipitation, Biacore, and MST studies confirmed specific binding between MIF and CXCL4L1 and determined a high-affinity binding constant in the 100-150 nM range for the interaction. Analysis of the binding interface by peptide array- based mapping and molecular docking provides an initial prediction of the residues involved in the CXCL4L1/MIF binding site.

### MIF/CXCL4L1 heterocomplex formation attenuates MIF-mediated inflammatory/athero- genic activities

We next wished to determine a potential functional role of MIF/CXCL4L1 heterocomplex formation. CXCL4L1 is a potent angiostatic chemokine acting through CXCR3 (Struyf *et al*, 2011), but its role in inflammatory responses and atherogenesis is not well understood. Pro- atherogenic activities of MIF have been extensively characterized and are mainly mediated through non-cognate interaction of MIF with CXCR2 and CXCR4 (Bernhagen *et al*., 2007; Sinitski *et al*., 2019). Here, we hypothesized that MIF/CXCL4L1 complex formation could predominantly influence CXCR4-mediated pathways of MIF.

We first asked whether MIF-elicited T-cell chemotaxis, a well-characterized atherogenic MIF effect mediated via T-cell-expressed CXCR4 (Bernhagen *et al*., 2007), is affected by CXCL4L1. Scouting experiments using Jurkat T-cells confirmed that, when added to the lower chamber of a Transwell migration device as a chemotattractant, MIF elicited chemotaxis with a chemotactic index (CTX) of approximately 2. Moreover, when CXCL4L1 was preincubated with MIF to allow for complex formation, no upregulation of Jurkat T-cell chemotaxis was observed, while CXCL4L1 alone exhibited neither a chemotactic nor inhibitory effect (*Supplementary Figure 3A*). In line with the observed lack of binding between MIF and CXCL4, MIF-mediated Jurkat T-cell chemotaxis was not attenuated by co- incubation with CXCL4, which by itself did not significantly enhance Jurkat T-cell chemotaxis (*Supplementary Figure 3B*). To test the physiological relevance of this finding, we next studied primary CD4^+^ T-cell chemotaxis and also applied a three-dimensional migration set-up, following individual cell migration trajectories by live cell imaging. MIF potently triggered T-cell migration as evidenced by a significant increase in forward migration index (FMI) (*Figure 3A-B*), confirming previous data showing CXCR4-dependent stimulation of monocyte migration by MIF (Kontos *et al*., 2020). This effect was abrogated when MIF was coincubated with CXCL4L1, while CXCL4L1 alone had no effect on 3D T-cell motility. This suggested that MIF/CXCL4L1 heterocomplex formation interferes with MIF/CXCR4-stimulated chemotaxis of T cells.

**Figure 3.**
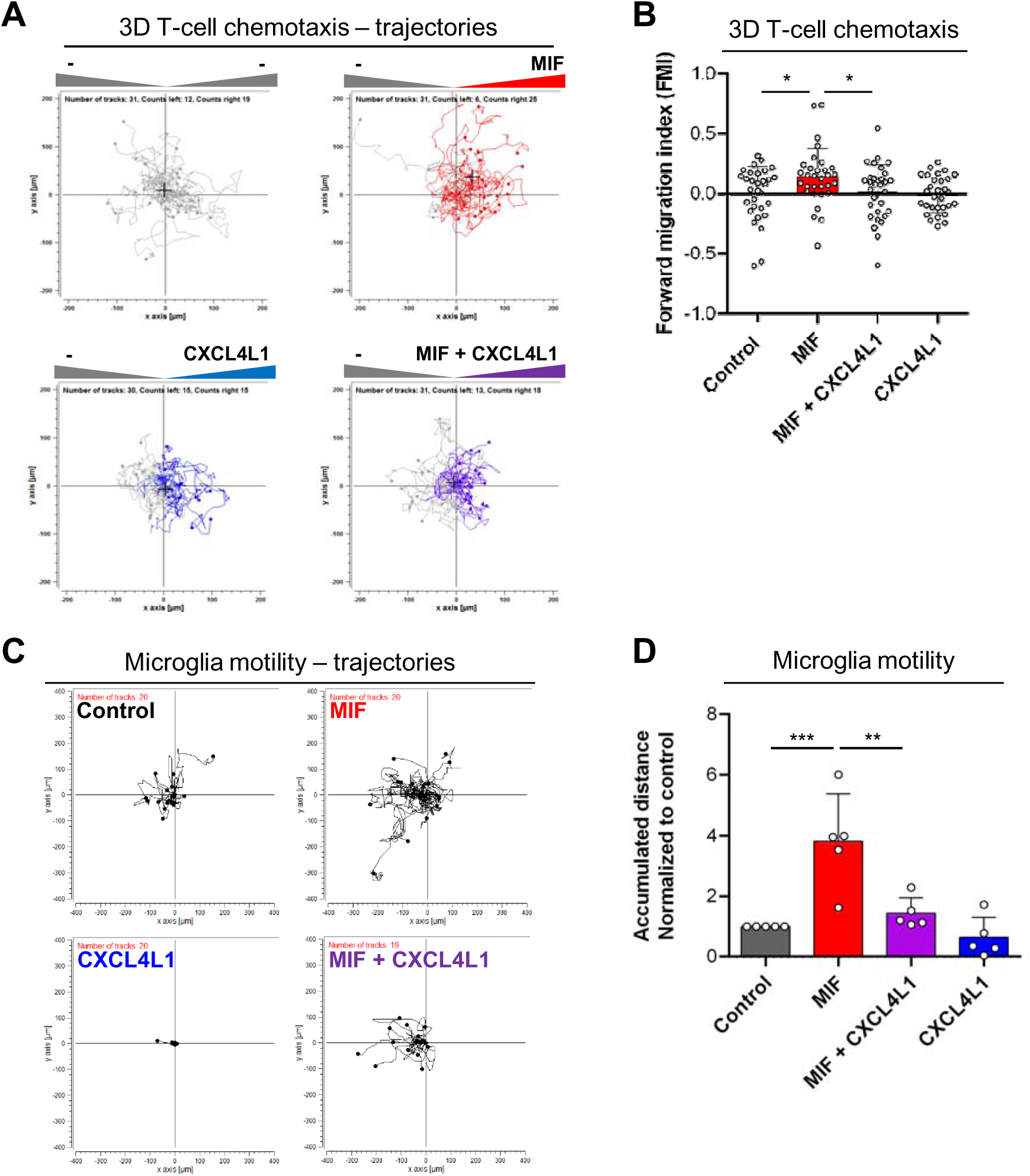
Co-incubation with CXCL4L1 inhibits MIF-mediated immune cell chemotaxis. (**A**) Migration of human CD4^+^ T-cells embedded in a gel matrix, subjected to gradients of MIF, CXCL4L1 or both. Movement of cells was followed by live cell imaging and individual tracks reconstructed from acquired images. Tracks of cells migrating towards the indicated stimuli are marked in the corresponding color. Starting point was centered to x = y = 0. The black crosshair indicates the cell population’s center of mass after migration. (**B**) Quantification of the 3D chemotaxis experiment in (**A**), indicating that complexation of MIF by CXCL4L1 attenuates MIF-mediated directed migration of human CD4^+^ T-cells. Plotted is the calculated forward migration index (FMI), based on manual tracking of at least 30 individual cells per treatment. (**C**) Migration trajectories of murine microglia, obtained by live cell imaging for 15 h, treated with MIF, CXCL4L1, or both. Used concentrations: MIF: 8 nM, CXCL4L1: 1.6 nM; n=5 independent experiments; horizontal bar: 100 µm. (**D**) Analysis of microglia motility, based on each tracked cell accumulated distance, shown in (**C**). Data is presented as mean ± SD. Statistical significance is indicated as described: *, P < 0.05; **, P < 0.01; ***, P < 0.001.

To study the potential relevance of these findings for other inflammatory/immune cell types, we next evaluated the effect of MIF/CXCL4L1 complex formation on microglial motility in the physiological setting of cortical brain cultures. MIF promotes the motility of Egfp^+^ microglia in murine cortical brain cultures *ex vivo* in a Cxcr4-dependent manner, as read out by live microscopy and as indicated by blockade of the MIF effect by the soluble CXCR4 mimicking peptide msR4M-L1(*Supplementary Figure 3C*). Importantly, MIF-triggered microglia migration in this setting was fully ablated when CXCL4L1 was added together with MIF following preincubation, while CXCL4L1 alone had no effect on microglia motility (*Figure 3C-D*). This indicated that CXCL4L1/MIF heterocomplex formation attenuates MIF’s CXCR4- dependent effect on microglia migration.

### MIF/CXCL4L1 heterocomplex formation inhibits MIF binding to CXCR4

The cell migration experiments implied, but did not directly test, the notion that MIF/CXCL4L1 complex formation affects MIF signaling through the CXCR4 pathway. To test the involvement of CXCR4 directly, we performed a binding competition experiment that capitalized on our recent identification of a MIF-binding CXCR4 ectodomain-mimicking peptide msR4M-L1 (Kontos *et al*., 2020). Employing fluorescence polarization spectroscopy (FP), titration of increasing concentrations of msR4M-L1 with Alexa 488-MIF led to a pronounced sigmoidal change in the FP signal (*Figure 4A*), in line with previous data showing high affinity binding between MIF and msR4M-L1 (Kontos *et al*., 2020). By contrast, when Alexa 488-MIF was preincubated with CXCL4L1 before the titration, the FP signal was ablated (*Figure 4A*), suggesting that MIF/CXCL4L1 heterocomplex formation interfered with MIF binding to the CXCR4 mimic.

**Figure 4.**
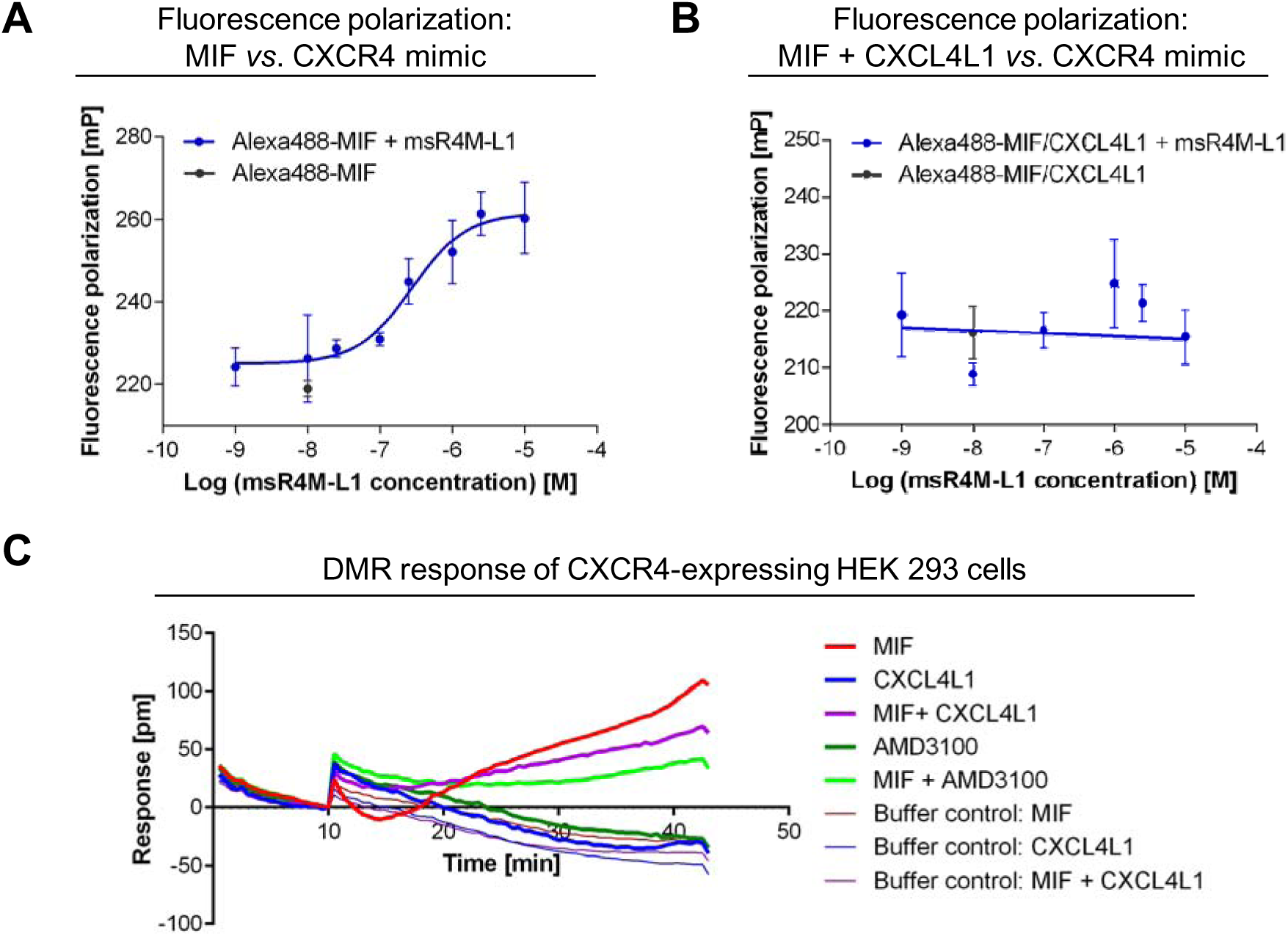
MIF/CXCL4L1 complex formation inhibits binding of MIF to CXCR4 and signaling of MIF through the CXCR4 signaling axis. (**A**) Fluorescence polarization (FP) spectroscopy shows the interaction of Alexa488-labeled MIF with the soluble CXCR4 receptor mimic msR4M-L1 with an apparent K_D_ of 237.2 ± 24.2 nM. Data is presented as mean of 3 independent experiments; error bars represent the SD. (**B**) Pre-incubation of MIF with CXCL4L1 (160-fold molar excess) prevents the interaction of MIF with msR4M-L1 (app. K_D_ > 10 µM). Mean of 3 experiments ± SD. (**C**) Dynamic mass redistribution (DMR) measurements with HEK293 cells stably expressing CXCR4 indicate that the cellular response to MIF is reduced, when MIF is pre-incubated with CXCL4L1. The DMR response of CXCR4-expressing HEK293 cells to MIF in the presence or absence of the CXCR4- antagonist AMD3100 is also shown, confirming the CXCR4-dependency of the cellular response to MIF.

To further confirm an interference of heterocomplex formation with the MIF/CXCR4 pathway, we next studied dynamic mass redistribution (DMR) responses in HEK293 cells stably transfected with human CXCR4. Incubation of HEK293-CXCR4 transfectants with MIF but not control buffer led to a pronounced time-dependent increase in the DMR signal as a real-time readout of an integrated cellular response of living HEK293 cell activation through the MIF/CXCR4 receptor signaling pathway (*Figure 4C)*. This signal was markedly attenuated by the small molecule CXCR4 inhibitor AMD3100, whereas the DMR curve of AMD3100 alone was similar to the control buffer curve, confirming CXCR4-dependency of the MIF-induced signal. Of note, preincubation of MIF with CXCL4L1 led to an appreciable reduction in the DMR response curve as well, when compared to cell stimulation with MIF alone, while CXCL4L1 alone and buffer control showed no effect (*Figure 4C)*.

Together, the competition binding study and the DMR experiment confirmed the notion that complexation by CXCL4L1 interferes with binding of MIF to CXCR4 and its ability to activate CXCR4-mediated cell responses.

### MIF and CXCL4L1 colocalize and form complexes in human platelet aggregates and clinical thrombus specimens

CXCL4L1 is an abundant platelet chemokine (Karshovska *et al*, 2013; von Hundelshausen *et al*, 2007) and we previously found that platelets also are a rich source of MIF (Strüßmann *et al*, 2013). The colocalization of MIF and CXCL4L1 in sub-cellular platelet compartments has not yet been studied, but a cell biological characterization of CXCL4 suggested that this paralog may be localized in a different intracellular platelet compartment than MIF (Strüßmann *et al*., 2013). Notwithstanding, we surmised that colocalization and complex formation between MIF and CXCL4L1 may occur extracellularly after secretion from activated platelets.

Initial evidence for a colocalization of CXCL4L1 and MIF following co-secretion from activated platelets came from human platelet preparations that aggregated due to handling stress. Examination of these aggregates by multi-photon microscopy (MPM) using an Alexa 488 signal to label MIF and Cy3 immunofluorescence for CXCL4L1 revealed several areas with an apparent colocalization of MIF and CXCL4L1 (*Figure 5A*). Colocalization was also detectable in areas with more isolated non-aggregated platelets (*Figure 5B*). These areas were then subjected to an in-depth analysis by fluorescence lifetime imaging-Förster resonance energy transfer (FLIM-FRET) capitalizing on the Alexa 488/Cy3 FRET donor-/acceptor pair. For molecule-molecule interactions within a distance range of 1-10 nm, FLIM- FRET monitors the change in fluorescence lifetime of the donor via FRET and directly visualizes the proximity of the donor (*here*: Alexa 488-labeled anti-mouse IgG secondary antibody in combination with mouse anti-MIF) and the acceptor molecule (*here*: Cy3-labelled anti-rabbit secondary antibody in combination with rabbit anti-CXCL4L1). We detected significant donor lifetime shortening (from 2.019 ± 0.069 ns to 1.496 ± 0.033 ns) and FRET events (FRET efficiency peak at 20-25%), when Alexa 488/Cy3 FLIM-FRET was recorded in appropriate regions-of-interest (ROIs) (*Figure 5C-D*), an observation that is consistent with the notion that MIF and CXCL4L1 not only colocalize in activated platelet preparations but form true heterocomplexes.

**Figure 5.**
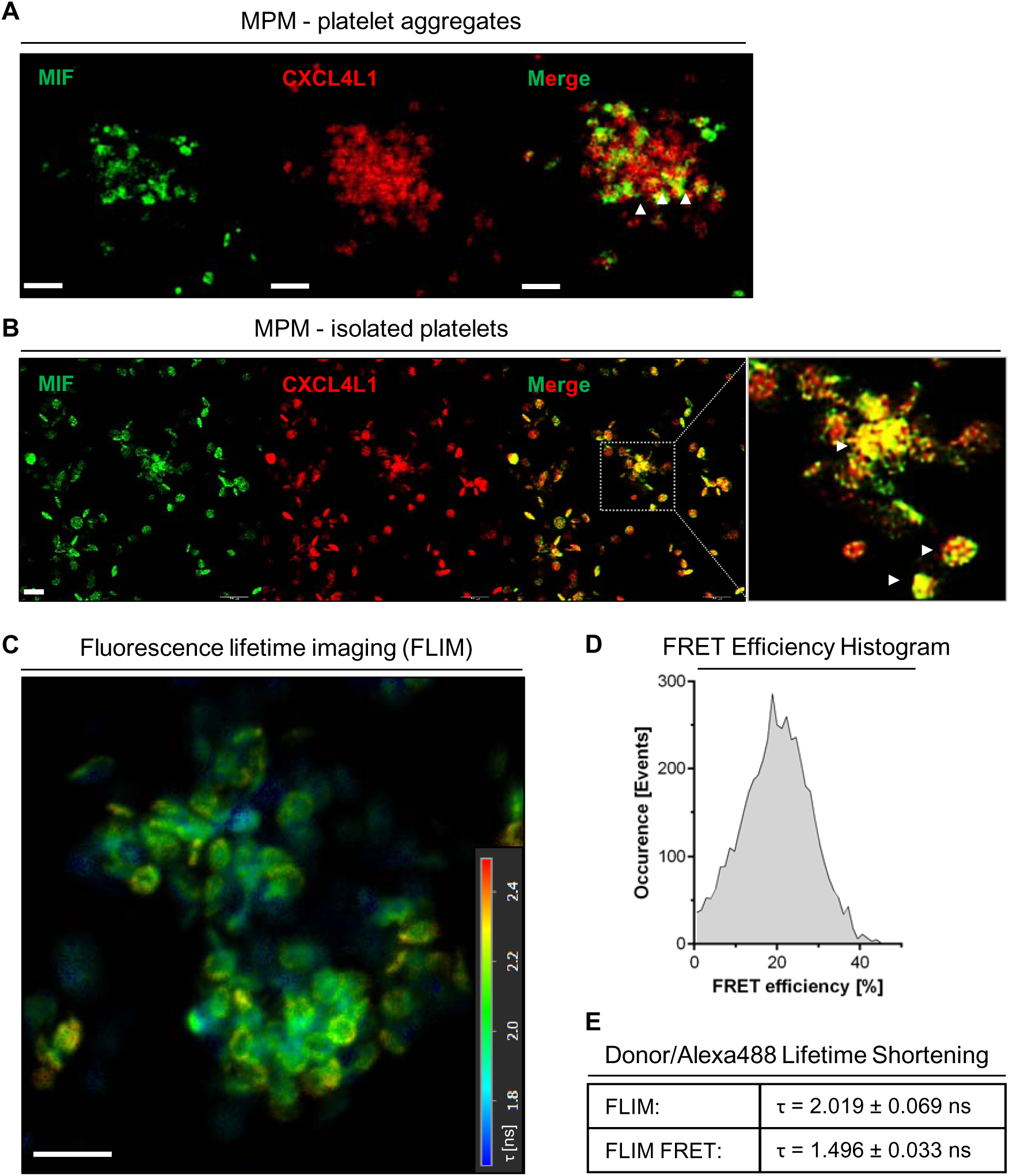
Co-localization and interaction of MIF and CXCL4L1 in human platelet preparations, detected in multiphoton microscopy (MPM). (**A**): MPM images of isolated platelets, forming small aggregates, stained for MIF and CXCL4L1. White arrowheads indicate areas of colocalization. Size bar: 5 µm. (**B**) MPM images of isolated, more separated platelets, stained as in (**A**), showing colocalization of MIF and CXCL4L1. Size bar: 5 µm. (**C**) Fluorescence lifetime imaging (FLIM) of platelets isolation as shown in (**B**). Color-code corresponds to lifetime of the donor, Alexa 488, the dye used for the antibody-based staining of MIF. (**D**) Histogram of the Förster Resonance Energy Transfer (FRET) efficiency in (**C**). (**E**) Donor lifetime shortening, presented as the mean lifetime (τ), average weighted, of the donor (Alexa 488, MIF staining) alone, and in combination with the acceptor fluorophore (Cy3, CXCL4L1 staining), where FRET occured.

To further investigate the physiological relevance of these findings, we next examined clinical thrombus specimens derived from vascular surgery procedures. To determine whether colocalized MIF and CXCL4L1 formed heterocomplexes in thrombus tissue, a proximity ligation assay (PLA) was performed which detects inter-molecular interactions within a distance of <10 nm. Specific PLA signals were detected in an atherosclerotic thrombus specimen (*Figure 6A-B*), suggesting the abundant occurrence of MIF/CXCL4L1 heterocomplexes in the context of clinical thrombus tissue and confirming the FLIM-FRET data obtained in platelet preparations from healthy blood samples. Thus, both FLIM-FRET and PLA demonstrated that MIF and CXCL4L1 form heteromeric complexes upon release from activated platelets.

**Figure 6.**
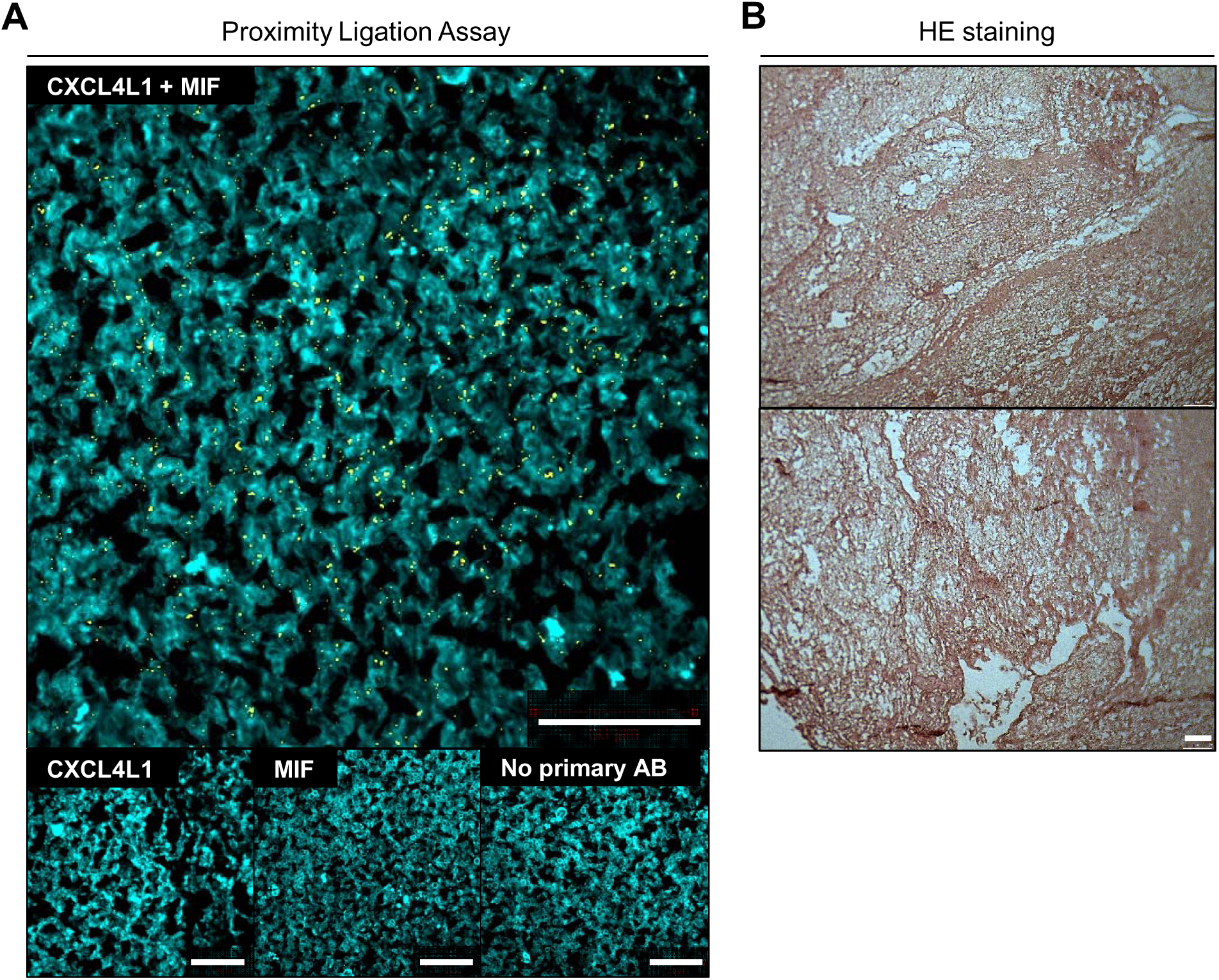
Proximity ligation assay (PLA) indicates that MIF/CXCL4L1 heterocomplexes are present in human thrombus tissue. (**A**) MIF/CXCL4L1 complex formation in thrombus specimen revealed by PLA. PLA-positive signals are depicted in yellow; tissue was counterstained with fluorescent-labeled phalloidin (cyan). Stained tissue samples were imaged by CLSM; size bar: 50 µm. (**B**) HE staining of thrombus tissue specimen; size bar: 75 µm.

### Heterocomplex formation inhibits MIF-stimulated thrombus formation and alters the effect of MIF on platelet morphology

Thrombus formation and clot retraction are relevant processes upon vessel injury and in advanced atherosclerotic vessels. MIF was found to modulate these processes (Wirtz *et al*, 2015). As our data showed that MIF/CXCL4L1 heterocomplexes form in the micro- environment of a thrombus, we next determined whether heterocomplex formation affects thrombus characteristics. Thrombus formation under flow perfusing diluted human blood over a collagen-coated surface harboring combinations of MIF and CXCL4L1 was studied as established (Chatterjee *et al*, 2014) and was found to double following exposure to MIF when applying a shear rate of 1000 s^−1^ (*Figure 7*). CXCL4L1 alone did not affect thrombus characteristics, but when added together with MIF following preincubation, MIF-elicited thrombus formation was blocked. These effects were mainly related to thrombus size/coverage (*Figure 7B*, *Supplementary Figure 5*) rather than thrombus numbers (*Figure 7C*). These data indicated that heterocomplex formation inhibited MIF-stimulated thrombus formation.

**Figure 7.**
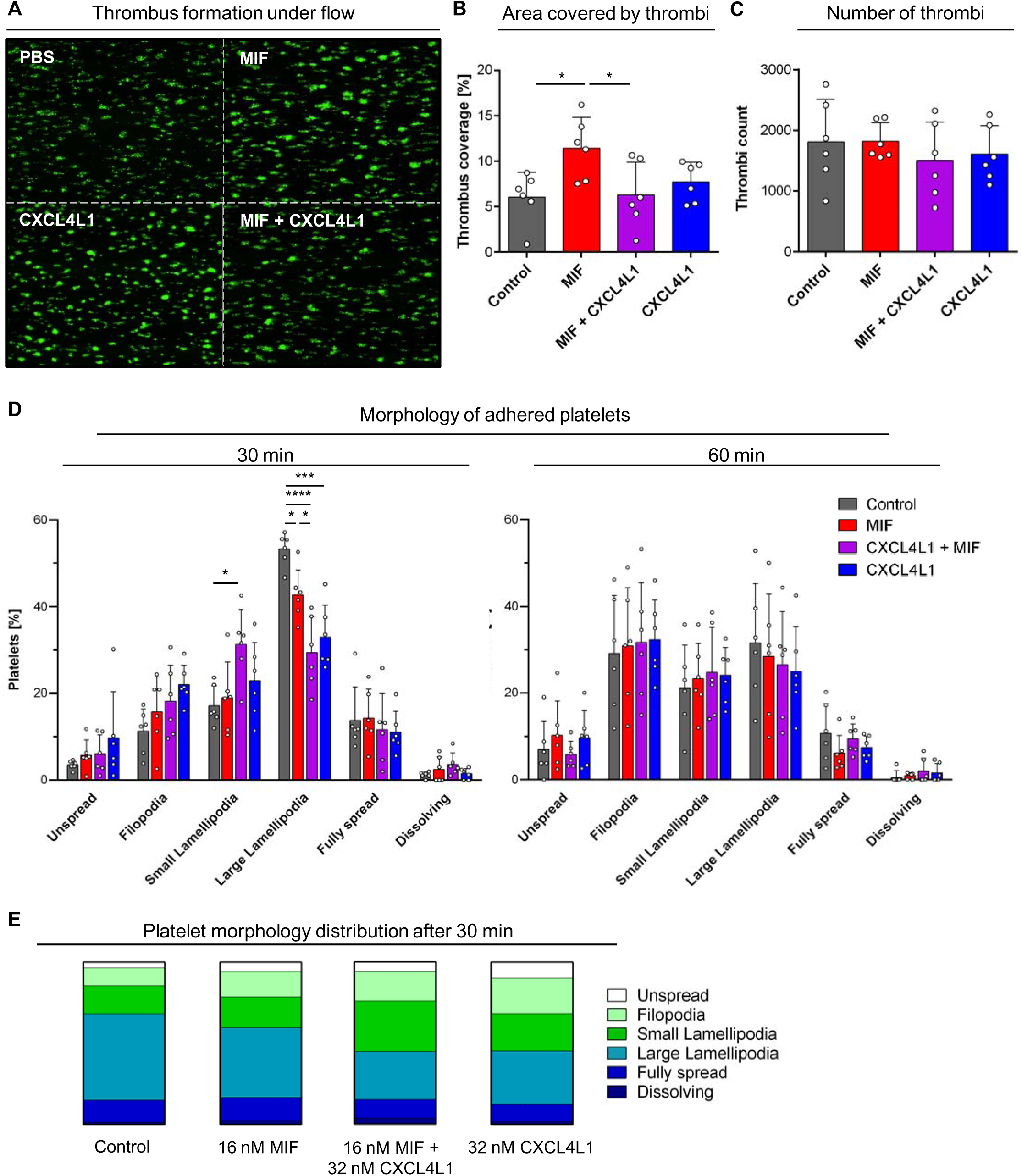
(**A**) Thrombus formation in human blood under flow stress is enhanced by MIF, and this effect is diminished by pre-incubation of MIF with CXCL4L1. Fluorescent staining with DiOC_6_. Shown are representative images of one experiment, performed at a shear rate of 1000 s^-1^; size bar: 100 µm. (**B**) Quantification of thrombi sizes from flow chamber experiments, as depicted exemplarily in (**A**). MIF-mediated increase in thrombus-covered area is diminished, when MIF is pre-incubated with CXCL4L1. n = 6 experiments and platelets coming from 4 donors. (**C**) Quantification of total thrombi numbers per treatment group. As thrombus numbers remain unchanged, effects on thrombus-covered area originate from the size of the formed thrombi (see also Supplementary Figure 5); n = 6 experiments. (**D**) Analysis and quantification of platelet morphology upon adhesion on fibrinogen-coated coverslips. Activated platelets were allowed to adhere on fibrinogen-coated coverslips that were pre-treated with MIF, CXCL4L1 or a mixture of both for the indicated times. After fixing with PFA, images of randomly selected areas were taken and platelet morphology analyzed. Treatment with a combination of MIF and CXCL4L1 led to a reduction in the large lamellopodia phenotype, favoring small lamellopodia, with the MIF/CXCL4L1 complex showing a stronger effect then the individual proteins; n = 6 experiments. (**E**) Platelet morphology distribution after 30 min for each treatment group according to panel (**D**).

The role of platelet morphology and lamellipodia in stable thrombus formation has been controversial, but platelet lamellipodia formation is critical for thrombus formation under flow (Fotinos *et al*, 2015; Kraemer *et al*, 2011a; Schurr *et al*, 2019). To further study the above observed effect of heterocomplex formation on thrombus behavior, we examined the morphology of flow-stressed adhered platelets exposed to MIF or heterocomplexes in detail. Platelet flow stress responses were recorded after 30 and 60 min, with significant changes observed for the 30 min time point. Morphological changes encompassed increased platelet numbers with filopodia, small lamellipodia, large lamellipodia, as well as fully spread platelets. Interestingly, the strong increase in large lamellipodia under control buffer conditions was significantly reduced by MIF and a further significant reduction was observed for platelets coincubated with MIF and CXCL4L1. Inversely, the incubation with the heterocomplex resulted in a significant increase in platelets with small lamellipodia compared to stimulation with MIF alone (*Figure 7D*). *Figure 7E* further illustrates the inverse effect of MIF/CXCL4L1 on large *versus* small lamellipodia formation. Together, these experiments indicated MIF/CXCL4L1 heterocomplex formation skewed the morphology of adhering flow- stressed platelets from a large to a small lamellipodia phenotype compared to treatment with MIF alone.

## Discussion

Chemokines control numerous pathogenic pathways contributing to inflammation and atherogenesis. The recent systematic characterization of the chemokine interactome revealed that heteromeric interactions between classical CC- and/or CXC-type chemokines represent an important molecular adjustment screw that serves to amplify, inhibit, or modulate chemokine activity (von Hundelshausen *et al*., 2017). Here, we have identified a heteromeric interaction between MIF, a pleiotropic inflammatory cytokine and ACK, and the classical platelet chemokine CXCL4L1. We also show that CXCL4L1/MIF complex formation affects inflammatory/atherogenic and thrombogenic activities of MIF. The scheme in *Figure 8* summarizes the main findings of this study. This suggests that disease-relevant activities of MIF may be fine-tuned by heterocomplexation with CXCL4L1 and that the chemokine interactome extends to heteromeric interactions between classical and atypical chemokines.

**Figure 8:**
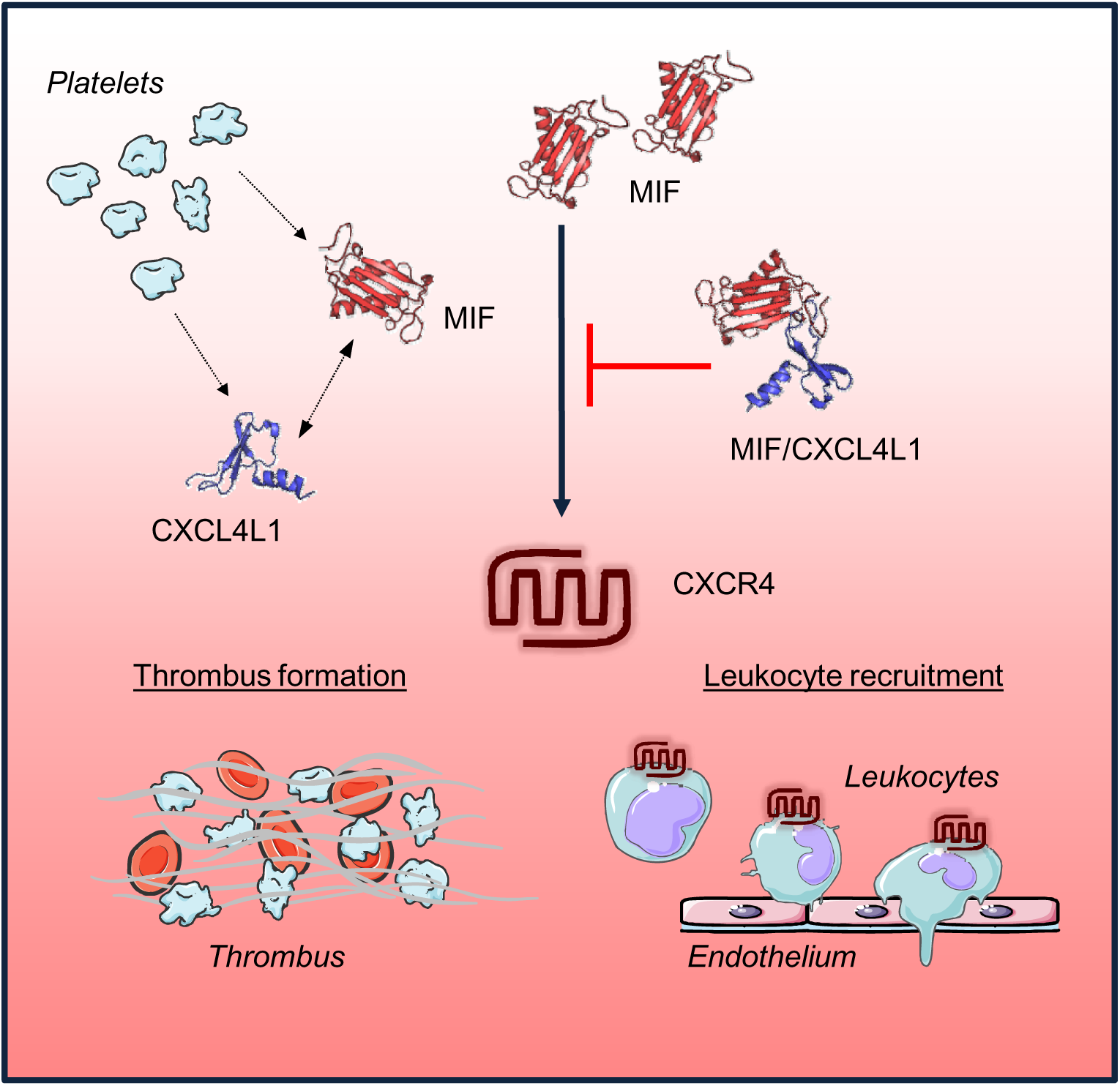
Summary scheme and suggested model of CXCL4L1/MIF complex formation and functions. The atypical chemokine MIF and the classical chemokine CXCL4L1, e.g. present in an inflammatory or atherogenic microenvironment after release from platelets, form heteromeric complexes. Complexes inhibit inflammatory effects of MIF on leukocyte recruitment as well as its pro-thrombotic effects through impairing MIF interactions with its non-cognate receptor CXCR4.

In fact, binding of classical chemokines to non-CC- or CXC-chemokine mediators is not unprecendented. Three examples have been documented: i) the CXC-chemokine CXCL12 binds to the alarmin HMGB1 and HMGB1/CXCL12 complex formation promotes chemotactic activity through CXCR4 (De Leo *et al*., 2019; Schiraldi *et al*., 2012); ii) the anti- microbial peptide and α-defensin HNP1 binds to CCL5 and enhances monocyte adhesion through CCR5 (Alard *et al*, 2015); iii) macrophage-expressed galectins such as galectin-3 (Gal-3) bind to CXCL12 and attenuate CXCL12-stimulated signaling via CXCR4 (Eckardt *et al*., 2020). However, while these studies underscore that classical chemokine activity may be modulated by interaction with various soluble mediators, HMGB1 and Gal-3 have no chemotactic activity on their own; HNP1 has been reported to exhibit chemoattractive properties, but the mediating chemoattractant receptor has remained elusive. In contrast, despite lacking the signature structural elements of classical chemokines such as the chemokine-fold and the N-terminal cysteine motif, MIF is a chemoattractant and depending on the microenvironmental context, can signal through the CXC chemokine receptors CXCR2, CXCR4, and/or ACKR3 to promote atherogenic and inflammatory leukocyte recruitment. Its CXC receptor binding capacity is based on the presence of a pseudo-ELR motif and an extended N-like loop, structurally mimicking the site 1 and 2 receptor binding motifs of the corresponding cognate ligands CXCL1/8 and CXCL12, respectively. Together with the β-defensins HDB1/2 and HBD3, which bind to CCR6 and CXCR4, respectively, and secreted fragments of certain AARSs, which bind to CXCR1 and CXCR2, MIF has therefore been designated an ACK (Degryse & de Virgilio, 2003; Kapurniotu *et al*., 2019; Oppenheim & Yang, 2005; Rohrl *et al*., 2010; Sinitski *et al*., 2019; Wakasugi & Schimmel, 1999). Our current identification of MIF/CXCL4L1 heterocomplexes thus also shows that the chemokine interactome is not strictly limited to interactions between classical CC- and/or CXC-type chemokines, but also encompasses heteromeric interactions between classical and atypical chemokines, with potential functional modulation of the chemokine receptor pathway of both the classical or atypical chemokine. Although not further validated and pursued in our current study, the detection of additional candidate interactors of MIF in our performed unbiased chemokine array, i.e. CCL28, CXCL9, as well as Prx6 leads us to hypothesize that interactions between classical and atypical chemokines could represent a broader principal of an “expanded ACK/CK interactome”.

The validity of the solid phase chemokine array as an unbiased screening approach for candidate chemokine interactors has been previously established (von Hundelshausen *et al*., 2017). The general utility and specificity of this methodology was further confirmed in the current study. Out of 47 immobilized classical chemokines, in addition to CXCL4L1, only two other classical chemokines, i.e. CCL28 and CXCL9, were revealed to have positivity. While a functional link between MIF and CCL28 has yet to be unveiled, it is interesting to note that the other detected CXC chemokine was CXCL9, a CXCR3 agonist like CXCL4L1. Intriguingly, biotin-MIF neither bound to CXCL12 nor to CXCL8, indicating that implicated functional interactions between MIF and the cognate CXCR4 and CXCR2 ligands, respectively, are independent of heterocomplex formation.

Futhermore, the specificity of the performed array is underscored by the notion that CXCL4, the highly homologous sister variant of CXCL4L1, did not bind to MIF, both at pH 8 and also when we tested for this interaction at pH 6 (data not shown) to account for pH-dependent charge differences. We hypothesize that the striking difference between CXCL4L1 and CXCL4 in binding to MIF might be due to the suggested different conformation of these two chemokines, e.g. the more exposed and flexible α-helix of monomeric CXCL4L1 (Kuo *et al*., 2013). While CXCL4 has been amply characterized by us and others as a pro- atherogenic platelet chemokine, in part also via its intriguing capacity to hetero-oligomerize with CCL5 (Koenen *et al*., 2009; von Hundelshausen *et al*., 2007), very little is known about the role of CXCL4L1 in chronic inflammatory diseases and atherosclerosis. Like its sister molecule, CXCL4L1 is also abundantly expressed in platelets; however, it apparently is not localized in α-granules but resides in a different sub-cellular compartment, from where it is constitutively secreted (Lasagni *et al*, 2007). It is also found in other cell types including mononuclear cells and smooth muscle cells (Lasagni *et al*., 2007). CXCL4L1 serves as an inhibitor of angiogenesis and has pro-inflammatory effects by inducing the release of CCL2 and CXCL8 from monocytes, while – contrary to CXCL4 – it does not promote monocyte survival (Domschke & Gleissner, 2019; Gouwy *et al*, 2016; Sarabi *et al*, 2011). There is only one *in vivo* study, in which CXCL4L1 was investigated as prognostic marker in cardiovascular disease. Interestingly, below-median levels of CXCL4L1 were found to correlate with a worse outcome in stable coronary artery disease patients, as indicated by a higher rate of cardiac death, stroke, or myocardial infarction (De Sutter *et al*, 2012). This finding might argue for a beneficial role of this chemokine in cardiovascular disease, even though the mechanisms behind this remain unclear, but certainly more studies are required. Of note, there is no equivalent of CXCL4L1 in mice (Eisman *et al*, 1990), limiting functional *in vivo* studies of this chemokine and its complex with MIF, as predicted from our study.

Importantly, we validated the binding between MIF and CXCL4L1 by semi-endo- genous pulldown from monocytes, as well as two different biophysical *in vitro* methods, i.e. SPR and MST. The combination of both methods also addresses potential disadvantages of having one interaction partner immobilized (Zhou *et al*, 2016). The binding affinity constants derived from the SPR and MST experiments (116 and 160 nM, respectively) are in reasonable agreement with each other. The observed (small) difference could be due to a number of factors, including surface immobilization effects, fluorescence *versus* biotin labeling, or buffers employed. Together, the results are suggestive of a relatively high binding affinity between MIF and CXCL4L1. Moreover, the obtained nanomolar K_D_ is consistent with the reported concentrations of both proteins in inflammatory disease settings (Sinitski *et al*., 2019). Flanking evidence for MIF/CXCL4L1 complex formation was obtained by our peptide array mapping and molecular docking results. As expected given their high sequence identity, the peptide array predicted identical binding sites for CXCL4 and CXCL4L1. Also, the peptide array methodology interrogates linear binding epitopes but cannot delineate conformational differences. In fact, Kuo *et al*. suggested that the three-amino acid difference between CXCL4 and CXCL4L1, although marginal, leads to a slight tilting of the C-terminal α-helix (Kuo *et al*., 2013). We hypothesize that this moderate conformational change could be the basis for the observed preferred binding of MIF to CXCL4L1 compared to CXCL4. Differences in their binding affinity to CCL5 have already been reported for CXCL4 and CXCL4L1 and also the availability of their monomers, regulated by the stability of their tetrameric complexes, differs between these two chemokines (Sarabi *et al*., 2011). Future structural studies, e.g. by nuclear magnetic resonance (NMR) spectroscopy, may help to further address these and other conformational questions.

To investigate the functional consequences of MIF/CXCL4L1 heterocomplex formation, we focused on inflammatory and atherosclerosis-relevant activities of MIF. T-cell migration is one such activity that is regulated by the MIF/CXCR4 pathway (Bernhagen *et al*., 2007). In line with previous results, MIF promoted T-cell migration in a physiologically relevant 3D migration setting. Although T cells generally express the CXCL4L1 receptor CXCR3, CXCL4L1 alone had no effect on the chemotaxis of human PBMC-derived T cells. Lack of CXCL4L1 activity in this assay is likely due to the fact that CXCL4L1 is not a *bona fide* T-cell chemoattractant (Gouwy *et al*., 2016) and that the preferential CXCL4L1 receptor variant CXCR3B is poorly expressed on T cells (Korniejewska *et al*, 2011). The 3D T-cell migration data are supported by the result that MIF, but not the combination of MIF and CXCL4L1, promoted Jurkat T-cell migration in a 2D Transwell assay. Confirming the remarkable specificity of MIF binding to CXCL4L1 *versus* CXCL4, coincubation of MIF with CXCL4 did not result in reduced Jurkat T-cell migration. Of note, heterocomplex formation with MIF led to a complete blockade of MIF’s pro-migratory effect on primary T cells in the 3D migration setting. While *in vivo* T-cell recruitment studies were beyond the scope of our study, inhibition of MIF-mediated T-cell migration by CXCL4L1 complexation could potentially be relevant in atherosclerosis, where it might represent a feedback mechanism that could serve to dampen the atherogenic response. In fact, abundant CXCL4L1 levels may be released by activated platelets in an atherogenic microenvironment, where they could colocalize with endothelial-immobilized or monocyte-secreted MIF and infiltrating T cells. That complexation of MIF by CXCL4L1 can interfere with MIF’s chemoattractant activities was confirmed in a microglia assay, in which the motility of Egfp^+^ microglia in murine cortical brain cultures *ex vivo* was studied. In addition to representing an independent cell migration system, the data obtained from the microglia-containing cortical cultures further confirmed that complex formation interferes with MIF signaling through the CXCR4 pathway and underscored that the mechanism could be relevant in *in vivo*-like physiological tissue settings. That MIF/CXCL4L1 heterocomplex formation interferes with MIF signaling through CXCR4 was independently validated by biochemical experiments using FP spectroscopy and DMR analysis of HEK293-CXCR4 transfectants.

This identified interaction of MIF with CXCL4L1, supposedly resulting in local inhibition of MIF’s pro-inflammatory effects was especially interesting to us in the context of previous studies, in which we identified human and mouse platelets as an abundant source of MIF (Strüßmann *et al*., 2013; Wirtz *et al*., 2015). Here, we verified expression and localization of MIF in human platelets as well as in platelet-rich clinical thrombus tissue by confocal (CLSM) and multiphoton microscopy (MPM). As expected, these experiments also showed the abundant presence of CXCL4L1 in platelets and thrombi, and suggested the colocalization and/or complex formation of MIF and CXCL4L1 in the vicinity of platelets. Due to the optical resolution limits of the CLSM and MPM methods, true colocalization and the specific subcellular compartment could not be determined. Evidence for the presence of MIF/CXCL4L1 heteromers is suggested by PLA performed on cryosections of a human thrombus. In fact, PLA is an established method to detect CK heteromers as shown previously for HNP1/CCL5 complexes (Alard *et al*., 2015).

Having confirmed the occurrence of this novel complex in platelet preparations and thrombus tissue, lastly the effect of MIF, CXCL4L1 and their complex on platelet function and thrombus formation was assessed. MIF promoted thrombus formation leading to a larger thrombus-covered area in an *in vitro* setting under flow conditions. Confirming our previous results, this effect was abrogated upon co-incubation with CXCL4L1. It is interesting to note that in the settings used in our experiment applying a shear rate of 1000 s^−1^ for 5 min, MIF acted to enhance thrombus formation. Instead, in a previous study employing a shear rate of 1700 s^−1^ MIF was found to reduce thrombus size, confirming that MIF is a modulator of thrombus formation, but also indicating that the directionality of the effect may depend on the specific microenvironmental context.

Moreover, studying the morphology change of isolated platelets during adhesion and activation on a fibrinogen-coated surface revealed that both MIF and CXCL4L1 favored a switch from large to small lamellipodia at an early time point. Interestingly, in this setting no inhibition by the complex on MIF-mediated effects was observed, but a synergistic behavior of MIF and CXCL4L1 was observed, suggesting that this effect may occur independently of CXCR4.

In addition to their classical role in wound closure and haemostasis, thrombus formation and platelet activation are processes that are closely linked to inflammatory processes driving atherosclerosis (Gawaz, 2006; Lippi *et al*, 2011; Nording *et al*, 2020; von Hundelshausen & Weber, 2007). MIF has been amply linked to atherosclerotic pathogenesis both clinically and experimentally, with evidence for a number of contributing mechanisms including leukocyte recruitment and platelet activation (Bernhagen *et al*., 2007; Chatterjee *et al*., 2014; Muller *et al*, 2013; Sinitski *et al*., 2019; Zernecke *et al*, 2008) The identified heteromerization of MIF and CXCL4L1 in our current study and the observed effect of MIF/CXCL4L1 complex formation on immune cell migration as well as thrombus size and platelet morphology might imply that CXCL4L1 could have a protective role in atherosclerosis by mitigating the pro-atherosclerotic effects of MIF via complex formation. This hypothesis would warrant future studies in corresponding experimental *in vivo* models, albeit the lack of CXCL4L1 expression in rodents will impose a particular challenge here.

In summary, we provide evidence that MIF does not only behave as a chemokine-like mediator by way of engaging classical chemokine receptors but also by direct binding to classical chemokines. Interestingly, the identified chemokine interactor of MIF is not one of the cognate ligands of the MIF receptors CXCR2 or CXCR4, but CXCL4L1, a prominent platelet chemokine not previously implicated in MIF biology or MIF-mediated pathologies. While evidence from experimental *in vivo* disease models will have to be obtained in future studies, our data suggest that MIF/CXCL4L1 complex formation could serve to attenuate inflammatory/atherogenic activities of MIF through the CXCR4 receptor axis. Our study also gives insight into the growing “chemokine interactome” with a particular focus on ACKs. While modulatory effects on the interactome by mediators not belonging to the class of chemokines have already been exemplified by intriguing studies involving HMGB1, HNP1, and the galectins (Alard *et al*., 2015; Eckardt *et al*., 2020; Schiraldi *et al*., 2012), the current study is first in demonstrating a role for MIF family proteins in particular, and *bona fide* ACKs in general, as defined by their chemotactic activity mediated through engagement of classical chemokine receptors. While not yet validated by follow up analyses, the identification of additional potential interactors in our array indicates that this could represent a broader principle of an ACK/CK interactome.

## Acknowledgements

This work was supported by Deutsche Forschungsgemeinschaft (DFG) grant SFB1123-A3 to J.B. and A.K., DFG INST 409/209-1 FUGG to J.B., SFB1123-A2 to P.v.H., SFB1123-A1 to C.W., SFB1123-B5 to L.M., SFB240-B01 (374031971 – TRR 240) to M.G., and by DFG under Germany’s Excellence Strategy within the framework of the Munich Cluster for Systems Neurology (EXC 2145 SyNergy—ID 390857198) to J.B., C.W., and O.G. A.H. was supported by a Metiphys scholarship of LMU Munich and by the Friedrich-Baur-Foundation e.V. at LMU University Hospital. E.S. and J.B. received funding from the LMU-FöFoLe program under project 52114016. C.W. is Van de Laar Professor of Atherosclerosis. We thank Simona Gerra, Priscila Bourilhon, and Lusine Saroyan for technical support, and Christine Krammer for help with the DMR assay. We are grateful to the mouse core facility of the Center for Stroke and Dementia Research (CSD) for their support. We also thank the Biophysics Core Facility at the School of Biology of LMU Munich and Dr. Sophie Brameyer for usage of the MST instrument.

## Supplementary figure legends

**Supplementary Figure 1.**
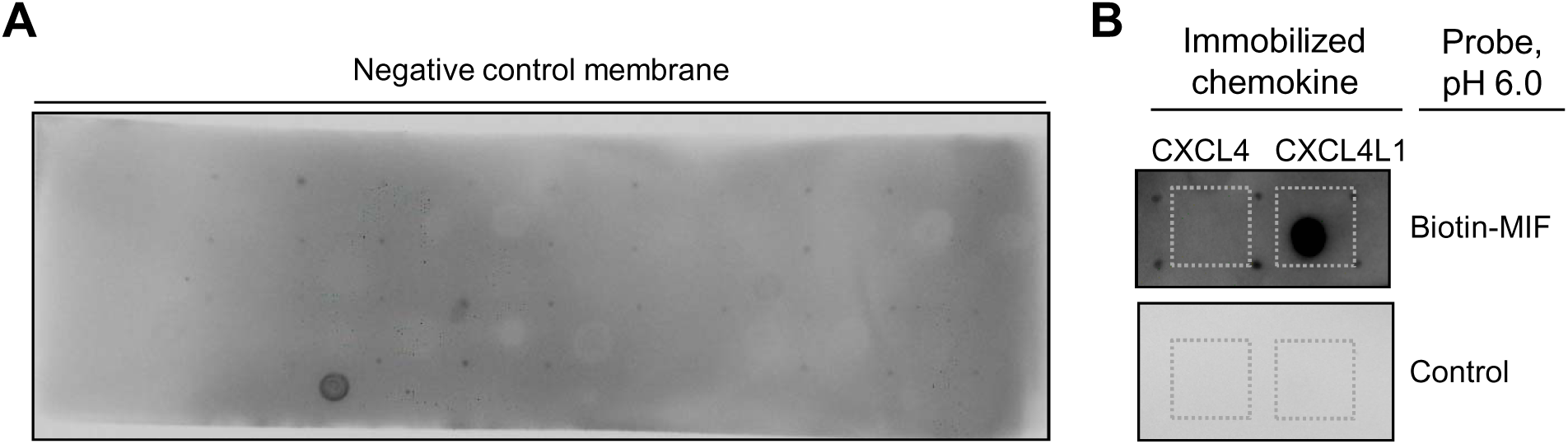
Additional data for chemokine protein array. (**A**) Negative control membrane related to the experiment in Figure 1, incubated in buffer at pH 8.0 without biotin-MIF. (**B**) Close-up of membrane from a chemokine protein array experiment with a focus on CXCL4 and CXCL4L1. The membrane was incubated with biotin-MIF and the incubation was performed at pH 6.0.

**Supplementary Figure 2.**
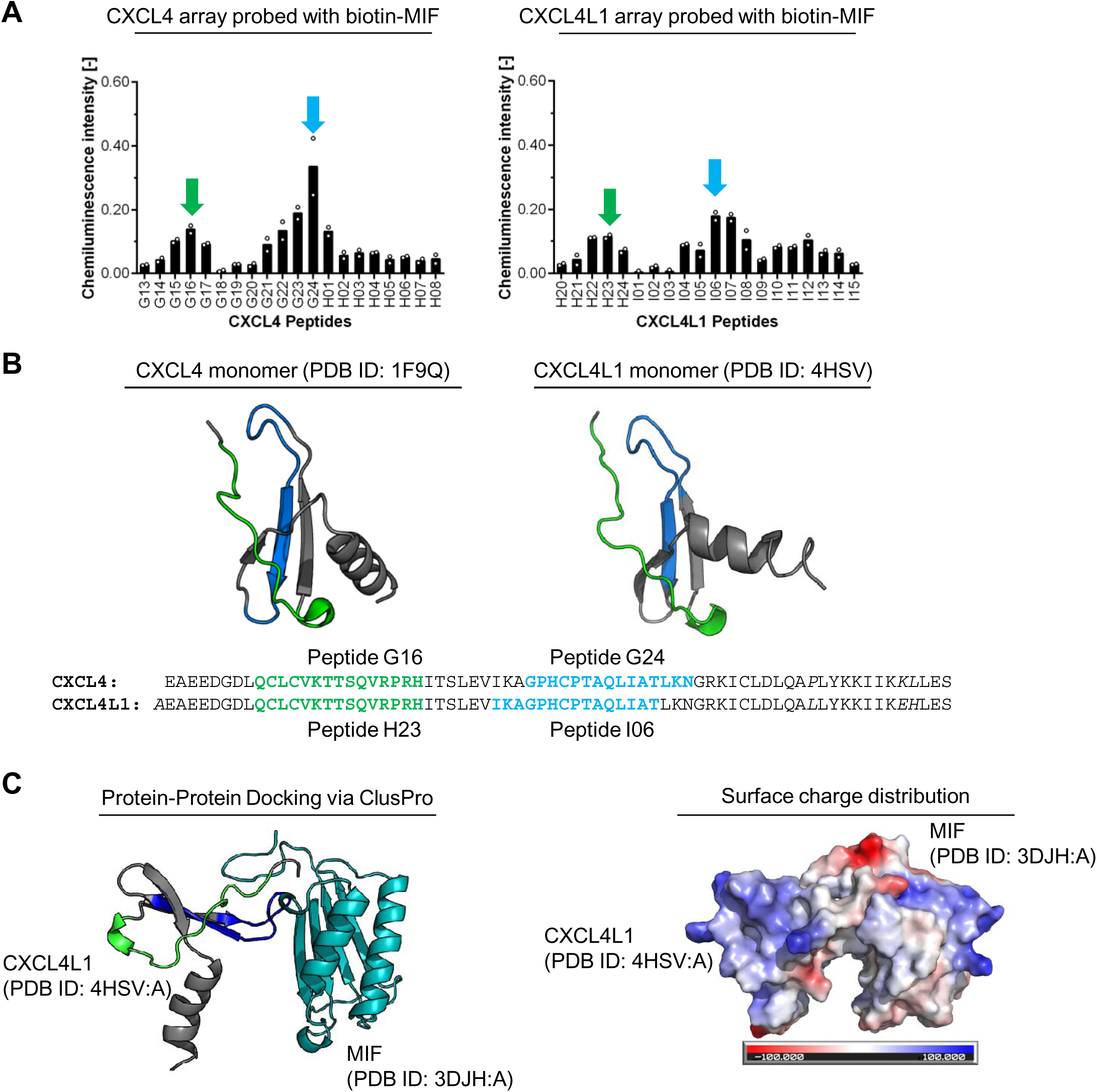
Investigation of the MIF/CXCL4L1 interaction interface and *in silico* studies. (**A**) CelluSpot peptide array experiments, where overlapping peptides of CXCL4 (*left*) and CXCL4L1 (*right*) were spotted on an array and probed with biotin-MIF. Chemiluminescence signal intensity indicates binding of biotin-MIF to the respective peptide. Arrows indicate peptides of interest that are most likely to be involved in the interaction with MIF. (**B**) Sequences of peptides identified in **A** are highlighted in the 3D structure of monomeric CXCL4 and CXCL4L1, showing their localization in the folded proteins. For both chemokines, these peptides of interest represent almost identical amino acid sequences, corresponding to highly similar regions of the protein. This indicates that not only the sequence but also the three-dimensional conformation of the chemokines might play a role in the interaction with MIF. Amino acid residues, in which CXCL4L1 differs from CXCL4 are in italics. PyMOL was used to visualize a CXCL4 (PDB ID: 1F9Q Chain A) and CXCL4L1 monomer (PDB ID: 4HSV Chain A). (**C**) To visualize the proposed MIF/CXCL4L1 complex, an unbiased *in silico* protein-protein docking approach was taken. The ClusPro 2.0 webserver was used to simulate a complex consisting of both a MIF and CXCL4L1 monomer. Depicted here is the highest-ranking docking result, with peptides identified in **A** to be potentially part of the interaction interface highlighted in CXCL4L1. According to this *in silico* prediction, they are partially directed towards MIF, allowing parts of their sequences being involved in complex formation. PyMOL was used to calculate the surface charge distribution of these proteins (red: negatively charged; blue: positively charged), revealing an area of opposite charges in the proposed contact region of MIF and CXCL4L1 that partially matches the peptide array results.

**Supplementary Figure 3.**
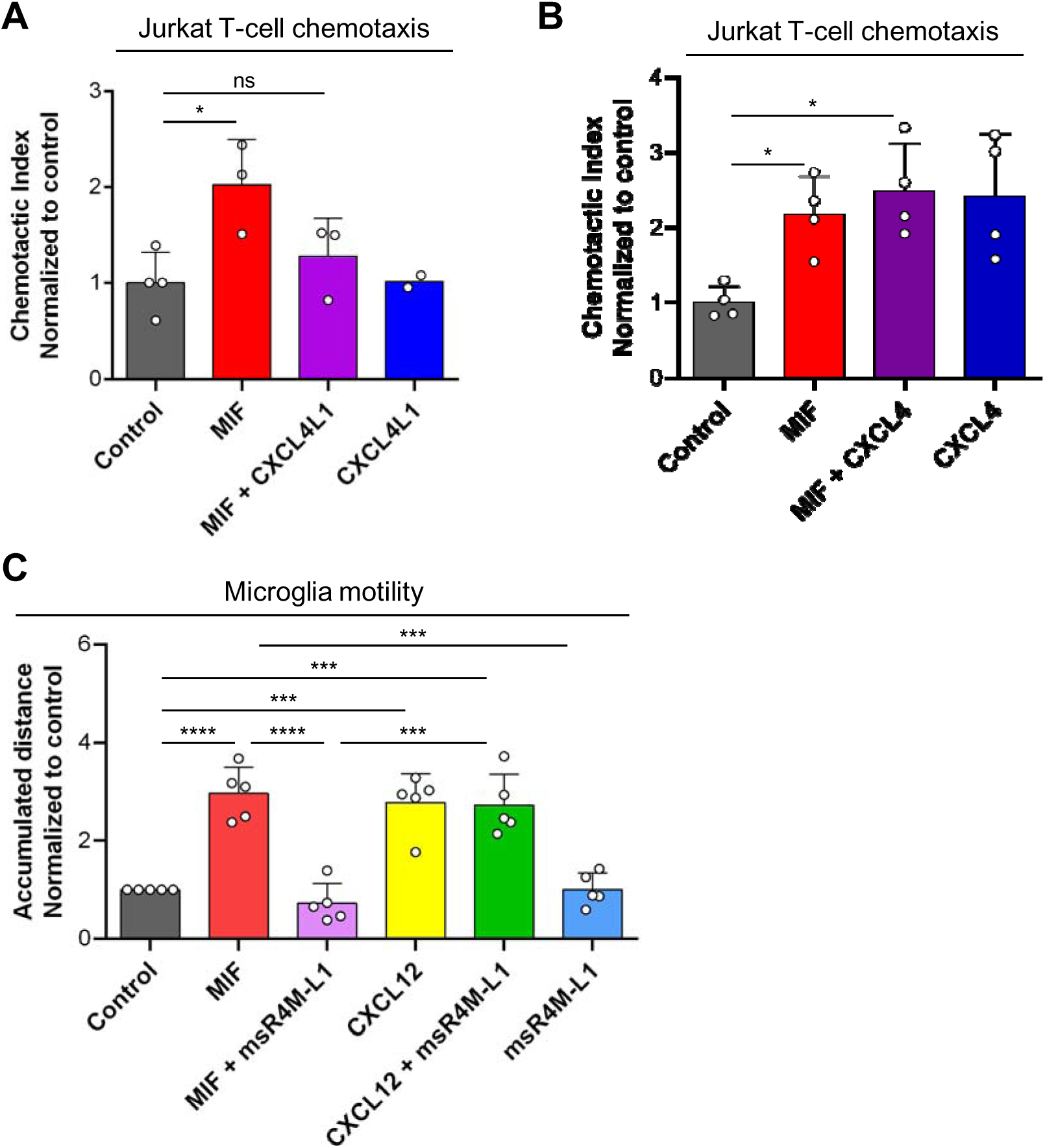
Effects on cell migration in Jurkat T cells and microglia. (**A**) Effect of CXCL4L1 on MIF-mediated chemotaxis of Jurkat T cells as analyzed in a Transwell migration assay. Used concentrations: MIF: 16 nM, CXCL4L1: 32 nM; Data is presented as mean ± SD. n = 2-4 independent experiments. (**B**) Same as (**A**), except that co-incubation with CXCL4 was analyzed. Data is presented as mean ± SD. n = 4 independent experiments with duplicates each. (**C**) Quantification of murine microglia motility, based on the accumulated distance of GFP-positive microglia tracked during live cell imaging (n = 5). MIF was used at a concentration of 8 nM, the soluble CXCR4-mimicking peptide msR4M-L1 at 40 nM and the cognate ligand of CXCR4, CXCL12, at 16 nM. Data presented as mean ± SD. Statistical significance: *, P < 0.05; **, P < 0.01; ***, P < 0.005; ****, P < 0.0001.

**Supplementary Figure 4.**
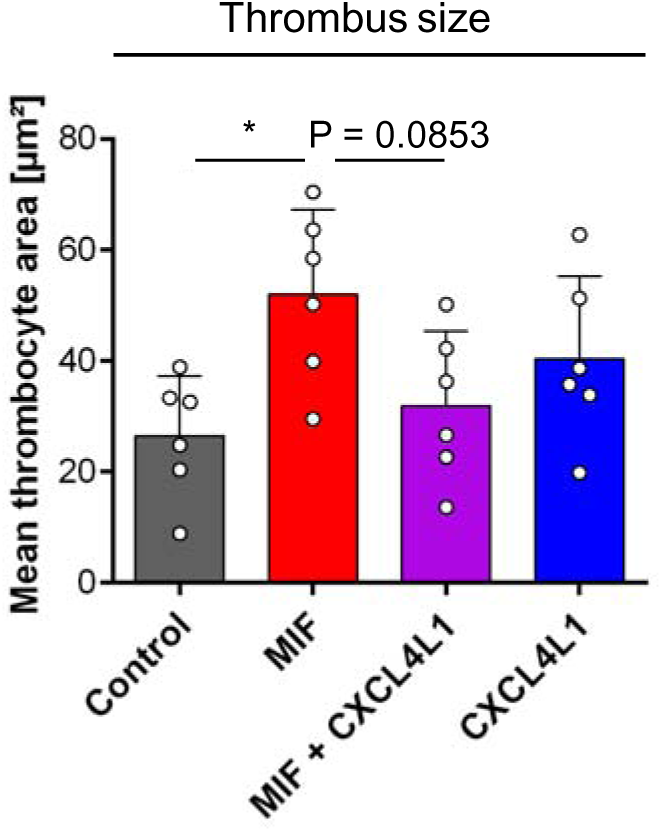
Quantification of mean thrombus sizes from Figure 5A, showing a trend for CXCL4L1 inhibiting the MIF-mediated increase in thrombus size in samples, in which MIF and CXCL4L1 were pre-incubated together.

